# Commanding or being a simple intermediary: how does it affect moral behavior and related brain mechanisms?

**DOI:** 10.1101/2021.12.10.472075

**Authors:** Emilie A. Caspar, Kalliopi Ioumpa, Irene Arnaldo, Lorenzo Di Angelis, Valeria Gazzola, Christian Keysers

## Abstract

History has shown that fractioning operations between several individuals along a hierarchical chain allows diffusing responsibility between components of the chain, which has the potential to disinhibit antisocial actions. Here, we present two studies, one using fMRI (Study 1) and one using EEG (Study 2), designed to help understand how commanding or being in an intermediary position impacts the sense of agency and empathy for pain. In the age of military drones, we also explored whether commanding a human or robot agent influences these measures. This was done within a single behavioral paradigm in which participants could freely decide whether or not to send painful shocks to another participant in exchange for money. In Study 1, fMRI reveals that activation in social cognition and empathy-related brain regions was equally low when witnessing a victim receive a painful shock while participants were either commander or simple intermediary transmitting an order, compared to being the agent directly delivering the shock. In Study 2, results indicated that the sense of agency did not differ between commanders and intermediary, no matter if the executing agent was a robot or a human. However, we observed that the neural response over P3 was higher when the executing agent was a robot compared to a human. Source reconstruction of the EEG signal revealed that this effect was mediated by areas including the insula and ACC. Results are discussed regarding the interplay between the sense of agency and empathy for pain for decision-making.

## INTRODUCTION

Numerous historical examples have shown the power of fractioning operations across different individuals to facilitate atrocious acts of mass annihilation (Swaan, 2015). A common example of fractioning operations are hierarchical situations: A superior communicates a plan and a subordinate executes it. The superior then bears responsibility for the decision but is distanced from the outcomes, while the subordinate experiences authorship over the action but may experience reduced responsibility for its outcomes (Bandura, 2006). In many organizations, orders are embedded in an even longer chain of commands in which a given commander often merely relays on the orders received from a superior. Commanders can thus also be intermediaries; an aspect which diffuses the psychological responsibility for the decisions to inflict harm. Experimental research has shown that such intermediary positions increase obedience to orders to hurt someone in comparison with being the author of that action or being the person giving the orders (Kilham & Mann, 1974; Milgram, 1974). However, the neural mechanisms by which being in the intermediary position or being in the commanding position disinhibit harming others remains largely unknown and represent the main focus of the present paper. In addition, modern warfare increasingly replaces the human soldiers that were at the bottom of the hierarchical chain and ultimately caused the harm to the enemy with artificial agents (e.g. drones, missiles, robots) (Di Nucci, & Santoni de Sio, 2016). How this affects the experience of commanders remains poorly understood.

Despite the lack of experimental research on the neuro-cognitive processes associated with moral behavior for intermediaries and commanders, previous scientific literature on the position of subordinate (i.e. the agent) has brought some evidence that at least two processes could be involved in how hierarchy influences the willingness to harm: sense of agency and empathy for pain.

The sense of agency refers to the feeling that we are the authors of, and thus potentially responsible for, our actions and their consequences in the external world (S. Gallagher, 2000). It is often measured implicitly through the intentional binding effect (Moore & Haggard, 2010): participants have to estimate the duration of the time interval between an action (e.g. pressing a button) and its consequences (e.g. hearing the beep it produces), with cases in which participants experience a stronger sense of agency leading to shorter time estimates (Moore & Haggard, 2010). The relationship between time perception and sense of agency is thought to be mediated by striatal dopaminergic activity, which is crucial for time perception (e.g.Meck, 2006) and is also driving information from basal ganglia to frontal motor areas (e.g. Nachev, Kennard, & Husain, 2008), key brain regions in generating the sense of agency. The feeling of responsibility is a related, but more explicit and social concept (Balconi, 2010), commonly evaluated with explicit questions asked to the participants (e.g. Li et al., 2011). Previous studies have shown that being in a position of the subordinate (or ‘agent’) executing an action commanded by an experimenter or induced by a computer reduces the sense of agency and the feeling of being responsible for an action (Caspar et al., 2016, 2018; Caspar, Ioumpa, et al., 2020; Barlas & Obhi, 2013). Other studies have also shown that asking participants to remember situations in which they had a low social power reduced their sense of agency compared to when they had a high social power (Obhi et al., 2012). It thus suggests that the sense of agency is reduced when people have a reduced power in social situations.

Empathy for pain is a fundamental process that allows us to understand and imagine what others feel by processing their pain within our own pain system. An extensive literature has indeed shown that seeing another individual in pain triggers an empathic response in the brain of the observer (Keysers & Gazzola, 2014; Decety, 2011; Lamm et al., 2019; Bernhardt & Singer, 2012). Empathy can be measured through several measurements, from subjective reports to neuroimaging measurements. In *f*MRI studies, the anterior cingulate cortex (ACC) and the anterior Insula (AI) have been shown to activate both when one experiences painful stimulations and when one empathizes with the same pain delivered to others (e.g. Jauniaux et al., 2019; Lamm et al., 2011; Timmers et al., 2018). In EEG studies, early (EAC, N200 - reflecting bottom-up emotional sharing response) and late potentials (LPP - representing a subsequent top-down evaluative response; Chen et al., 2014) are sensitive to the witnessing of a painful stimulus being delivered to another individual. Similarly to the sense of agency, being in a position of subordinate executing an action commanded by an experimenter reduces the empathic response to the pain of others (Caspar, Ioumpa et al., 2020; Lepron et al., 2015). Another study has shown that being reminded about a situation in which individuals had high social power increased their empathic neural response to painful pictures compared to being reminded about a low social power condition (Galang et al., 2021). These studies do suggest that being in a position with low decisional power impacts also empathy for pain. Finally, Cui et al., (2015) showed that if the participant’s action is not the only cause for a victim’s pain, neural signatures of empathy are reduced when witnessing the pain of the victim compared to cases in which the participant’s actions is the only cause for that pain.

However, these studies never compared directly the positions of commanding or being a simple intermediary in a single paradigm, thus preventing direct comparisons between those two social positions (Caspar et al., 2016, 2018; Caspar, Ioumpa, et al., 2020; Lepron et al., 2015). Further, in some of those studies, participant’s actions or decisions were not measured and participants were simple observers (Galang et al., 2021; Obhi et al., 2012). The present study aimed to fill this gap by comparing how commanding or being in an intermediary position impacts the sense of agency and empathy for pain within a single behavioral paradigm in which participants could decide or had to follow the orders to send or not painful shocks to another participant in exchange for money.

Participants were recruited in pairs and respectively played the role of the person giving orders or the role of the ‘victim’. When they were in the role of the person giving orders, participants had to give an order to an agent to send or not to send a real, mildly painful electric shock to the ‘victim’ in exchange for a small monetary gain, which increased their own remuneration for their participation in the study. In that position, participants were either free to decide which order to send to the agent (i.e., they were ‘commanders’) or were given an order by the experimenter that they had to transmit to the agent (i.e., they were ‘intermediaries’). When participants were in the commander position, we also modulated the entity executing their orders: they were either giving orders to another human or to a robot. As robots are considered entities with no personal responsibility, we wanted to understand if giving orders to a robot in the intermediary position would boost the sense of agency compared to giving orders to another human being, on which they can diffuse their own responsibility (Bandura et al., 1999).

In a first Study (Study 1), we used *f*MRI to investigate how the processing of the pain felt by the ‘victim’ for each shock received is modulated by the different experimental conditions by quantifying BOLD signals in regions associated with empathy while the participant witnessed the shock being delivered vs. not being delivered. In a second Study (Study 2), we used electroencephalography to further explore how the different experimental conditions modulated the pain processing as measured by the amplitude of the N2, P3 and late positive potentials (Coll, 2018), and the sense of agency, as measured by intentional binding effects on time interval estimation. The sense of agency was not measured using time interval estimation in Study 1 because, to separate brain activity related to motor response from those related to processing the pain of the victim in *f*MRI, long action-outcome intervals (i.e. between 2.5 and 6 s) have to be used (e.g., Caspar, Ioumpa, et al., 2020; Cui et al., 2015), which are too long for the measurement of the sense of agency with the method of interval estimates. Previous studies indeed showed that if the consequence of an action occurs more than 4s after the action, modulations of the sense of agency no longer lead to measurable changes in time perception (Buehner & Humphreys, 2009). We therefore used electroencephalography in Study 2, which has a better temporal resolution than *f*MRI, to investigate the two targeted neuro-cognitive processes in a single paradigm.

Based on past literature (Galang et al., 2021; Obhi et al., 2012), we expected that being in a low social power position (i.e., intermediaries) would reduce empathy for pain and the sense of agency compared to a position of higher power (i.e., commanders). This may be even more the case when orders are transmitted to another human compared to a condition in which orders are transmitted to a robot, as responsibility can be diffused between two humans but less in the case of a human-robot interaction (Ciardo et al., 2020). In a former study, results indicated that in hierarchical situations, the sense of agency was reduced for both the commander and the agent executing orders (Caspar et al., 2018). In the present study, we did not use an experimental condition in which people would also be in the position of the ‘agent’, as it would have considerably increased the testing time. However, we compared the results from Study 1 with those from a recent *f*MRI study using a matching design to empathy for pain for participants playing the role of the subordinate (i.e., referred to as ‘agents’ in Caspar, Ioumpa, et al., 2020). This comparison allowed us to understand how empathy for the pain of the ‘victim’ is modulated through three different hierarchical positions: commander, intermediary and agent. We also further conducted exploratory analyses in order to investigate how self-reported personality traits influenced prosocial behaviors.

## METHOD – STUDY 1 (MRI)

### Participants

Forty participants were recruited in 20 dyads, based on the number of participants recruited in Caspar, Ioumpa et al. (2020). None of the participants reported to know each other. The full dataset of three participants was excluded from all the analyses, two because of disobeying by contradiction (performing the reverse action from what they were ordered), and one because systematically disobeyed orders by administering shocks even when requested not to do so. fMRI data from one participant was removed from the fMRI analyses due to extensive movements, one because of scanner failure, and twelve because they delivered less than 5 shocks in each condition, causing a too little number of repetitions to measure reliable signals. An extra criterion was for participants to report correctly, through a pain scale, whether or not a shock was delivered on the victim’s hand (see *procedure and material* section below). No participants were excluded for this reason. These resulted in 23 participants for the fMRI (6 males; 24.26y±3.17) and 38 for the behavioral analyses (13 males; 25.05 y±3.6SD). The study was approved by the local ethics committee of the University of Amsterdam (project number: 2017-EXT-8298). Data is made available on OSF (*will be made public after publication*).

### Procedure and material

Upon arrival in the laboratory, both participants received instructions about the experiment and provided informed consent together, ensuring that they were each aware of the other’s consent. Then, their individual pain threshold for the electrical stimulation was determined, as described in Caspar et al. (2016). Two electrodes were placed on the participants’ left hand on the abductor pollicis muscle in order to produce a clear and visible muscle twitch and the threshold was increased by steps of 1mA until a mildly painful stimulation was achieved. The pain threshold was determined by asking a series of questions to the participants about their pain perception during the calibration (1. « Is it uncomfortable? » - 2. « Is it painful? » - 3. « Could we increase the threshold? »).

Participants were assigned to start either as ‘participant’ or ‘victim’ by randomly picking up a card in a box, but were offered the possibility to change if they wanted to. The participant who was in the role of the ‘commander’ was brought in the MRI scanner to perform the task, while the participant assigned to the role of the ‘victim’ was seated at a table in the nearby control room

‘Victims’ were asked to place their left hand on a black sheet positioned in the field of view of the camera and asked not to move their hand during the entire scanning session. The ‘victim’ was invited to watch a neutral documentary to make the time pass.

In order to be able to compare the MRI data acquired in this experiment with the MRI data from a previous study (Caspar, Ioumpa et al., 2020) in which participants played the role of the agents (i.e., agents), we preserved the exact same trial structure. Each trial started with a jittered fixation cross lasting 8-12 seconds, see **Fig. 1**. Then, participants heard a verbal instruction from the experimenter in all three experimental conditions. In the IntermediaryWithHumanAgent condition, the experimenter told participants to either ‘*give a shock*’ or ‘*don’t give a shock*’. To have a similar causation and auditory information in all the experimental conditions, participants also received a verbal instruction in the two Commander conditions. This verbal instruction was ‘*you can decide*’. Participants were told that the experimenter would give those instructions from outside the scanner room through the interphone during the two Commander conditions and from inside the scanner room during the IntermediaryWithHumanAgent condition. In reality, these sentences were pre-recorded in order to keep control on the precise timing of each event during the scanning session. To increase the authenticity of the procedure, each sentence was recorded 6 times with small variations in the voice to generate credible variance, and these recordings were presented in random order. In addition, the audio recordings included a background sound similar to interphone communications. Participants were also told that during the IntermediaryWithHumanAgent condition, the experimenter would wear a microphone to increase the intensity of her voice to overcome the noise of the MRI scanner. The experimenter explicitly exhibited herself at the beginning of the IntermediaryWithHumanAgent condition by speaking with the participant laying down inside the scanner, but then moved in the corner of the room to avoid visual interference due to her presence.

**Fig. 1.**
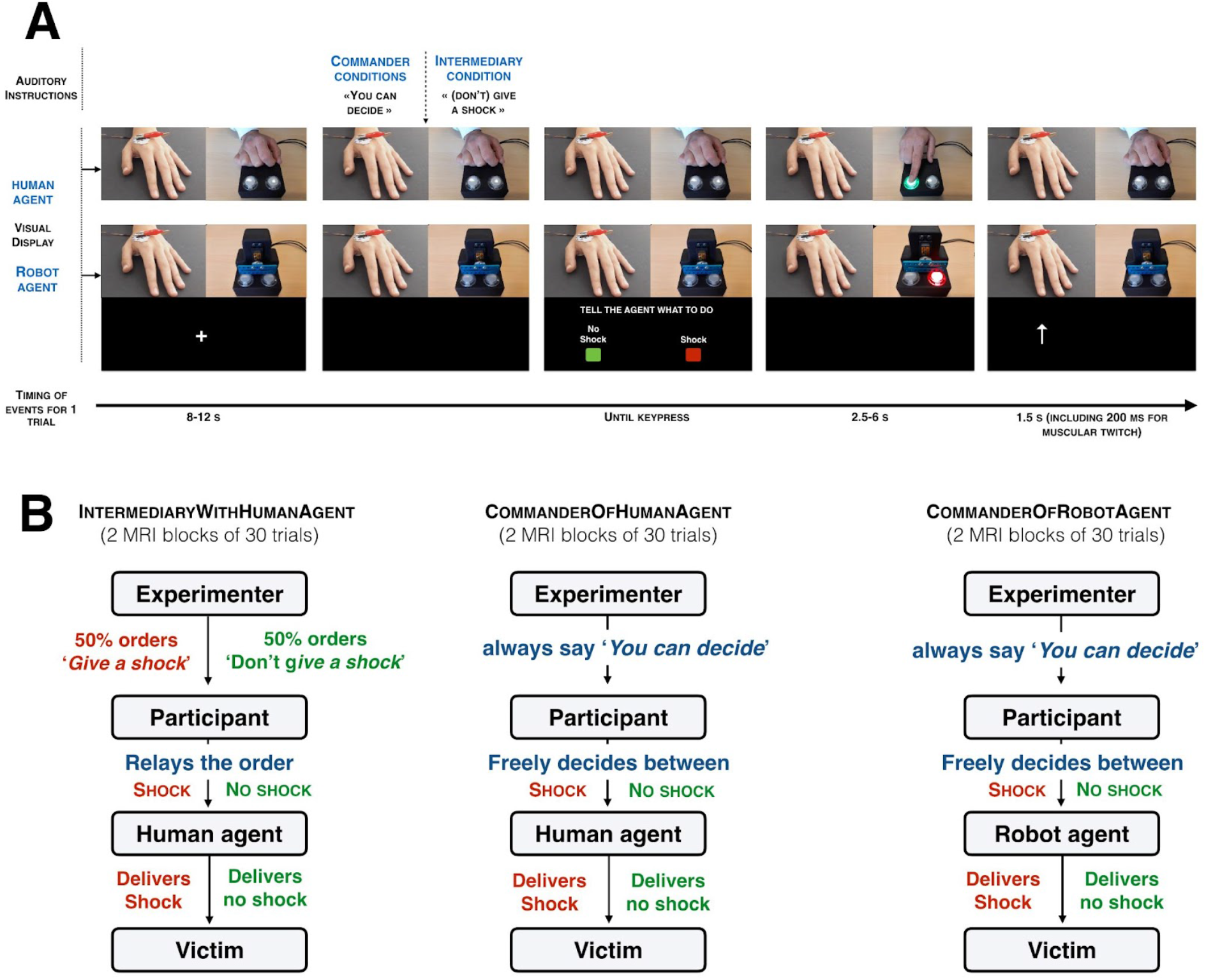
**(A) Visual display of the structure of a single trial.** Participants inside the MRI scanner had two real-time video feedbacks, one from the victim’s left hand with the electrodes connected to the shock machine and one with the agent (human or robot) and its button box. When the participants pressed either the “shock” or the “no shock” button, the corresponding buttons on the agent’s button box appeared in red or green. The agent then pressed on the colored button. An arrow pointing to the top then appeared on the screen to remind participants to look at the victim’s hand at that moment. If the “shock” button was pressed, participants could see a visible muscular twitch on the victim’s hand. **(B) Schematic representation of each experimental condition**. Each participant underwent 2 blocks of 30 trials in the scanner. In the two IntermediaryWithHumanAgent blocks, in half the trials they heard the experimenter tell them to ‘give a shock’, and then pressed a red button relaying this order to the human agent that they could then see press the corresponding red button and the victim’s hand then twiched. In the other half of the trials, the experimenter told them to ‘don’t give a shock’, and then had to relay that order by pressing the green ‘no shock’ button, leading the agent to press the green button as well. They could then see the victim not receiving a shock. In the four Commander blocks, they heard the experimenter tell them ‘you can decide’, and the participant in the scanner was then in a position of Commander, freely deciding whether to shock or not shock on each trial. In two of these blocks, the agent was again a human (CommaderOfHumanAgent). In the other two blocks, the human agent was replaced by a mechanical device that pressed the ordered button (CommanderOfRobotAgent)

After receiving the verbal order, a picture of two rectangles, a red one labeled ‘SHOCK’ and a green one labeled ‘NO SHOCK’, was displayed on the left and right bottom of the screen. The key-outcome mapping varied randomly across trials to concentrate motor preparatory activity in the interval between choice screen onset and key-press, but the outcome was always fully congruent with the participants decision, i.e. the agent never disobeyed the order given by the participant. Participants could then press one of the two buttons in order to ask the agent to execute their order. On the right top of the screen, participants could see the agent pressing a button corresponding to the order they had given. The agent had a button box with two transparent buttons. After commanders gave their order, the corresponding button popped in the corresponding color (red for SHOCK and green for NO SHOCK) so that the agent knew which button to press. This procedure ensured that participants could track the agent’s action. It also ensured that the agent was pressing the correct button corresponding to the requested order. Pressing the SHOCK button delivered a shock to the victim while pressing the NO SHOCK button did not deliver any shocks. The shock was delivered between 2.5 and 6 seconds after the keypress (Cui et al., 2015; Caspar, Ioumpa et al. 2020). For the participants to also track the consequence of their orders, another real-time camera feed displayed on the left top of the screen showed the victim’s hand, with electric shocks eliciting a visible muscle twitch. Seven hundred and fifty ms before the display of the shock, an arrow pointing to the top was displayed to remind participants to look at the video. This arrow also appeared when the NO SHOCK button had been pressed to keep a similar structure in all trials. That arrow disappeared 750 ms after. To ensure that participants were actually watching the victim’s hand, on 6 trials in each MRI run a pain rating scale appeared, ranging from ‘*not painful at all*’ (‘0’) to ‘*very painful*’ (‘1,000’). Participants were asked to rate the intensity of the shocks seen by moving the red marker bar along the scale using four buttons. The keys below the middle fingers allowed to modify the number associated with the position of the marker by steps of +-100. The keys below the index fingers allowed to modify the answer by steps of +-1. After a fixed duration of 6 seconds, their answer was saved and the next trial started. If no shocks were delivered on that trial, participants were asked to report that the shock was ‘*not painful at all*’.

In the IntermediaryWithHumanAgent, participants were asked to obey the experimenter’s orders and to transmit those orders to the agent. In the CommandFree condition, participants were entirely free to decide which order to send to the agent. In the CommanderOfHumanAgent condition, the agent was a human, confederate of the experimenter. Participants were told that the human agent was part of the experimenter’s team. In the CommanderOfRobotAgent condition, the agent was a robot. The experimenter confirmed that agents would always obey the commander’s order in all the experimental conditions.

The task was split into 6 blocks of 30 trials each, 2 blocks for the CommanderOfHumanAgent condition, 2 blocks for the CommanderOfRobotAgent condition and 2 blocks for the IntermediaryWithHumanAgent condition (presented in 6 separate *f*MRI acquisition runs). Order of the experimental conditions was counterbalanced across participants. Anatomical images were recorded between the fourth and fifth run of *f*MRI acquisition. At the end of each task block, participants rated their explicit sense of responsibility over the outcomes of their actions on an analogue scale presented on the screen, ranging from ‘*not responsible at all*’ to ‘*fully responsible*’. Each delivered shock was rewarded with +€0.05 in all the experimental conditions, and in IntermediaryWithHumanAgent blocks, participants were instructed to transmit an order to shock on 50% of trials.

At the end of the experimental session, participants were asked to fill out 8 questionnaires assessing several personality traits. Those questionnaires included (1) the Interpersonal Reactivity Index (IRI, Davis & Association, 1980), (2) the Short Dark Triad (DT, Jones, & Paulhus, 2014), (3) the Levenson Self-Report Psychopathy Scale (LSRP, Levenson et al., 1995), (4) the Moral Foundation Questionnaire (MFQ, Graham et al., 2011), (5) the Aggression-Submission-Conventionalism scale (ASC, Dunwoody & Funke, 2016), (6) the Right-Wing Authoritarianism scale (RWA, Altemeyer, 1981), (7) Hypomania Checklist (HCL, Angst et al., 2005), and (8) a debriefing assessing what they felt during the experiment (see ***Supplementary Material S1***). Participants were paid separately, based on their own gain during the experiment.

### General data analyses

Data were analyzed with both frequentist and Bayesian statistics (Dienes, 2011), except for voxel-wise brain analyses that were only analyzed using frequentist approaches. Bayesian statistics assess the likelihood of the data under both the null and the alternative hypothesis. In most cases, we report BF_10_, which corresponds to the *p*(data|*H*_1_)/*p*(data|*H*_0_). Generally, a BF between 1/3 and 3 indicates that the data is similarly likely under the H1 and H0, and that the data thus does not adjudicate which is more likely. A BF_10_ below 1/3 or above 3 is interpreted as supporting H_0_ and H_1_, respectively. For instance, BF_10_=20 would mean that the data are 20 times more likely under H_1_ than H_0_ providing very strong support for H_1_, while BF_10_=.05 would mean that the data are 20 times more likely under H_0_ than H_1_ providing very strong support for H_0_ (Marsman & Wagenmakers, 2017). BF and p values were calculated using JASP (Love et al., 2019, p. 2019) and the default priors implemented in JASP (Keysers et al., 2020). Default priors used in JASP depend on the statistical tests performed (for ANOVA, see Rouder et al., 2012; for t-tests, see Ly et al., 2016; for correlations, see Wagenmakers et al., 2016). In cases where a one-tailed hypothesis was tested, the directionality of the hypothesized effect is indicated as a subscript to the BF (e.g. BF_+0_ for a positive effect, BF_-0_ for a negative effect).

### Functional Magnetic Resonance Imaging (fMRI)

MRI images were recorded using a 3-Tesla Philips Ingenia CX system and a 32-channel head coil. T1-weighted structural images were recorded with the following specifications: matrix = 240×222; 170 slices; voxel size = 1×1×1mm. Six runs of functional images were recorded (matrix M x P: 80 × 78; 32 transversal slices in ascending order; TR = 1.7 seconds; TE = 27.6ms; flip angle: 72.90°; voxel size = 3×3×3mm, including a .349mm slice gap). Images were acquired in ascending order.

### General fMRI Data processing and first level contrasts

MRI data processing was carried out in SPM12 (Ashburner et al., 2014). EPI images were slice-time corrected to the middle slice and realigned to the mean EPI image. High quality *T*_1_ images were coregistered to the mean EPI image and segmented. The normalization parameters computed during the segmentation were used to normalize the gray matter segment (1mmx1mmx1mm) and the EPIs (2mmx2mmx2mm) to the MNI templates. Afterwards, images were smoothed with a 6mm kernel.

At the first level, we defined separate regressors for Shock and NoShock trials, with the three different conditions modeled in separate runs in order to identify the activations associated with witnessing pain. Each of these regressors started 750 ms before the moment of the shock, which lasted 250 ms, up to 500 ms after the moment of the shock. This moment corresponded to when the arrow pointing to the video feedback appeared, to remind participants to watch the screen displaying the victim’s hand. The same 1.500ms-time window was taken for Shock and NoShock trials. Additional regressors included: (1) The auditory orders from the experimenter (starting between 8-12s after the start of the trial) together with the button presses (participants could press the key whenever they wanted right after the auditory orders) and the presses of the (human or robot) agent, (2) the pain rating scale (appearing on 6/30 trials randomly 1s after the arrow pointing towards the video feedback disappeared) together with the responsibility rating scale (appearing at the end of each MRI run, 1s after the arrow pointing towards the video feedback disappeared or again 1s after the pain scale). Trials where participants disobeyed were modeled in additional regressors of no interest separately for ‘prosocial’ disobedience (i.e., they refused to administer a shock while having been ordered to send a shock) and ‘antisocial’ disobedience (i.e., they administered a shock while having been ordered not to send a shock). Finally, 6 additional regressors of no interest were included to model head translations and rotations.

At the first level, we defined three main contrasts of interest: [CommanderOfHumanAgent(S-NS)-IntermediaryWithHumanAgent(S-NS)],[CommanderOfRobotAgent(S-NS)-IntermediaryWithHumanAgent(S-NS)],[CommanderOfRobotAgent(S-NS)-CommanderOfHumanAgent(S-NS)].

We included the contrast of Shock-NoShock rather than examining the shock condition alone in each condition to isolate the effect of witnessing a shock from carry-over activity associated with pressing the response button and seeing the arrow presented during the feedback period. To be noted, some participants administered a few shocks. Because the reliability with which brain activity in the Shock condition can be estimated in the *f*MRI analysis depends on the number of trials included, only including participants delivering a large number of shocks would be ideal. However, for our results to be representative of the population, excluding too many participants delivering small numbers of shocks would bias our results towards less considerate participants.

At the second level, we tested whether the contrasts [CommanderOfHumanAgent(S-NS)-IntermediaryWithHumanAgent(S-NS)],[CommanderOfRobotAgent(S-NS)-IntermediaryWithHumanAgent(S-NS)],[CommanderOfRobotAgent(S-NS)-CommanderOfHumanAgent(S-NS)] were significant across participants in a random effect one-sample t-test.

## RESULTS - STUDY 1

### Number of shocks delivered

In the IntermediaryWithHumanAgent condition, participants (N=40) were ordered by the experimenter, on a trial-basis, to tell the agent to inflict 30/60 shocks to the ‘victim’, in random order. In the Commander conditions, participants could freely decide which order to send to the agentagent. In the CommanderOfHumanAgent condition, participants asked the human agent to administer 24.34/60 (SD=15.43, min: 0 – max: 59) shocks to the ‘victim’. In the CommanderOfRobotAgent condition, participants asked the robot agent to administer 24.13/60 (SD=15.13, min: 0 – max: 60) shocks to the ‘victim’. In the IntermediaryWithHumanAgent condition, the experimenter ordered to deliver shocks on 30 of the 60 trials. Of these, the participants relayed the shock order to the human agent on average 24.63/30 trials (SD=9.003, min: 0 – max: 30), whilst in the remaining trials, they disobeyed and ordered the agentagent not to deliver a shock. More specifically, out of the 40 participants, twelve reported that they voluntarily disobeyed the orders of the experimenter on some trials. Among those 12 participants, 10 disobeyed ‘prosocially’ by refusing to send a shock during ‘shock trials’ and by telling the agent not to deliver a shock to the ‘victim’ even if the experimenter asked them to do so (i.e. prosocial disobedience), and 2 disobeyed ‘by contradiction’, that is, they disobeyed as often on ‘don’t shock’ trials and on ‘shock trials’. We conducted a repeated-measures ANOVA with Condition (CommanderOfHumanAgent, CommanderOfRobotAgent, IntermediaryWithHumanAgent) as within subject factor and Role-Order (Commander first, Victim first) as between-subject factor on the number of shocks sent in each experimental condition. Results indicated that none of the main effects or their interactions were significant (all *p*>.3), see **Fig. 2A**. The Bayesian version of the same analysis indicated that the main effect of Condition and the interaction Condition × Order of the Role were strongly in favor of H_0_ (BF_incl_=.062 and BF_incl_=.033, respectively). The main effect of Order of the Role was slightly in favor of H_0_ (BF_incl_=.387).

**Fig. 2.**
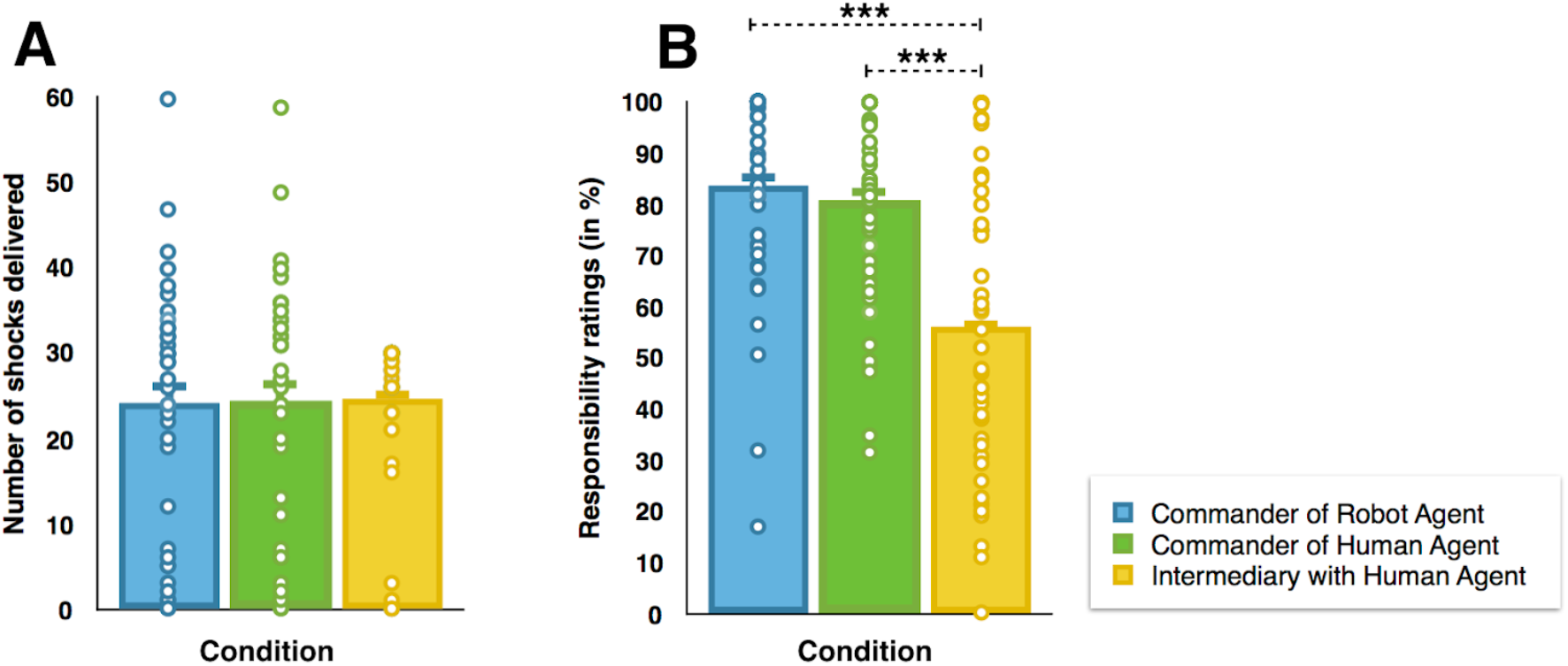
A) Graphical representation of the number of shocks delivered in the three experimental conditions. B) Graphical representation of responsibility ratings in the three experimental conditions. All tests were two-tailed. *** represents p <=.001 and a BF > 3.

### Responsibility ratings

At the end of each experimental condition, participants had to report how responsible they felt for the outcome of their orders. We conducted a repeated-measures ANOVA with Condition (CommanderOfHumanAgent, CommanderOfRobotAgent, IntermediaryWithHumanAgent) as within subject factor and Role-Order (Commander first, Victim first) as between-subject factor on the responsibility ratings. Both the frequentist and the Bayesian results supported a main effect of Condition (F_(2,74)_=25.038, *p* < .001, η^2^_***partial***_ = .404, BF_incl_=4.597E+6), see **Fig. 2B**. Paired-comparisons indicated that responsibility ratings were higher in the CommanderOfRobotAgent condition (82%, CI_95_=74.5-89.5) than in the IntermediaryWithHumanAgent condition (56.2%, CI_95_=46.8-65.6, t_(38)_=-5.455, *p* < .001, Cohen’s d=-.873, BF_10_=5822.75) and in the CommanderOfHumanAgent condition (81%, CI_95_=74.4-87.4) than in the IntermediaryWithHumanAgent condition (56.2%, CI_95_=46.8-65.6); t_(39)_=-5.050, *p* < .001, Cohen’s d=-.799, BF_10_=1889.49). The difference in responsibility ratings between the CommanderOfHumanAgent and the CommanderOfRobotAgent conditions was inconclusive (*p*>.1, BF_10_=.466). The main effect of Order of the Role (*p*=.063, BF_incl_=.968) and the interaction Condition × Order of the Role (*p*>.5, BF_incl_=.439) were inconclusive.

### Pain Scale

The estimations reported by agents on the pain scale were not analyzed as only 9/23 participants had data for shock trials in the three experimental conditions.

### fMRI whole brain analyses

We first ensured that we could detect the vicarious pain activation network in our study, including especially the anterior insula (AI) and the anterior cingulate cortex (ACC). We thus computed a main Shock-NoShock contrast, irrespective of the experimental condition. We observed typical pain observation network activation, including the ACC, MCC, SII and Insula (see **Fig. 3**) suggesting that witnessing the shock delivered on the victim’s hand indeed triggered an empathic neural response.

**Fig. 3 .**
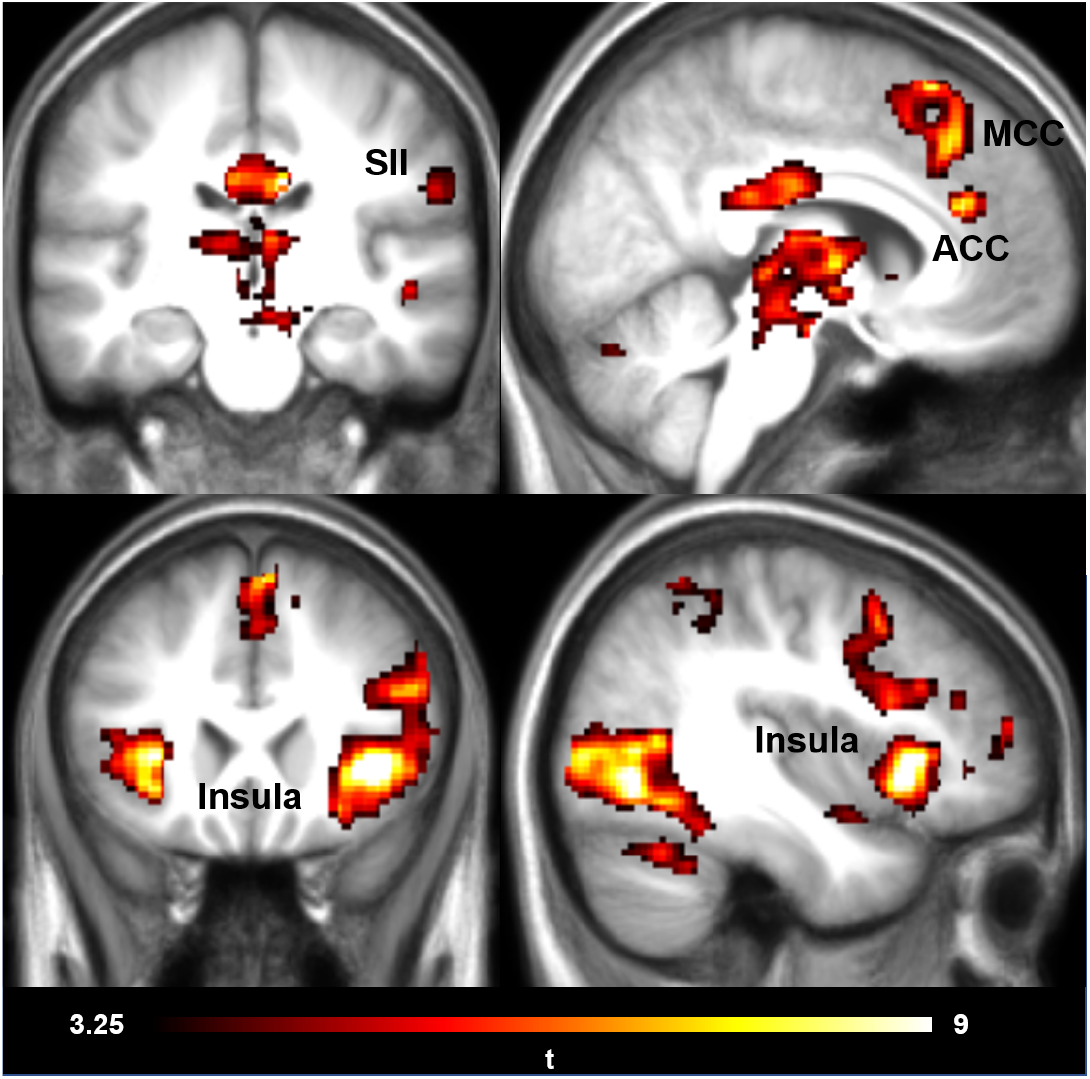
MRI results. Shocks - Noshock contrast for all experimental conditions together. Peak coordinates can be seen in Table S1. Results are shown thresholded using p_FWE_<0.05 at cluster level (cFWE=160 voxels) following a cluster cutting threshold at t=3.5, *p* < .001.

Results showed that at the whole brain level, none of our contrasts of interest [CommanderOfHumanAgent(S-NS)-IntermediaryWithHumanAgent(S-NS)],[CommanderOfRobotAgent(S-NS)-IntermediaryWithHumanAgent(S-NS)],[CommanderOfRobotAgent(S-NS)-CommanderOfHumanAgent(S-NS)] showed significant results in a random effect one-sample t-test.

### Comparison between being commander and agent

One of the aims of this study was to compare the empathic neural response when participants are in the role of a commander (MRI data acquired in the present experiment) and when they are in the role of an agent executing an order of a commander (MRI data from a previous study by Caspar, Ioumpa et al., 2020). At the second level, we thus conducted five two-sample t-tests comparing the two conditions of the previous agent study with the three conditions of the current commander study as follows:

[AgentFree(S-NS)-CommanderOfRobotAgent(S-NS)], [AgentFree(S-NS)-CommanderOfHumanAgent(S-NS)], [AgentFree(S-NS)-IntermediaryWithHumanAgent(S-NS)], [AgentCoerced(S-NS)-CommanderOfRobotAgent(S-NS)], [AgentCoerced(S-NS)-CommanderOfHumanAgent(S-NS)], [AgentCoerced(S-NS)-IntermediaryWithHumanAgent(S-NS)].

Results on **Fig. 4** were thresholded at punc < .001 and 5% family-wise error (FWE) corrected at the cluster level and significant activation was observed for the [AgentFree(S-NS)-IntermediaryWithHumanAgent(S-NS)], [AgentFree(S-NS)-CommanderOfHumanAgent(S-NS)] and [AgentCoerced(S-NS)-CommanderOfHumanAgent(S-NS)]

**Fig. 4:**
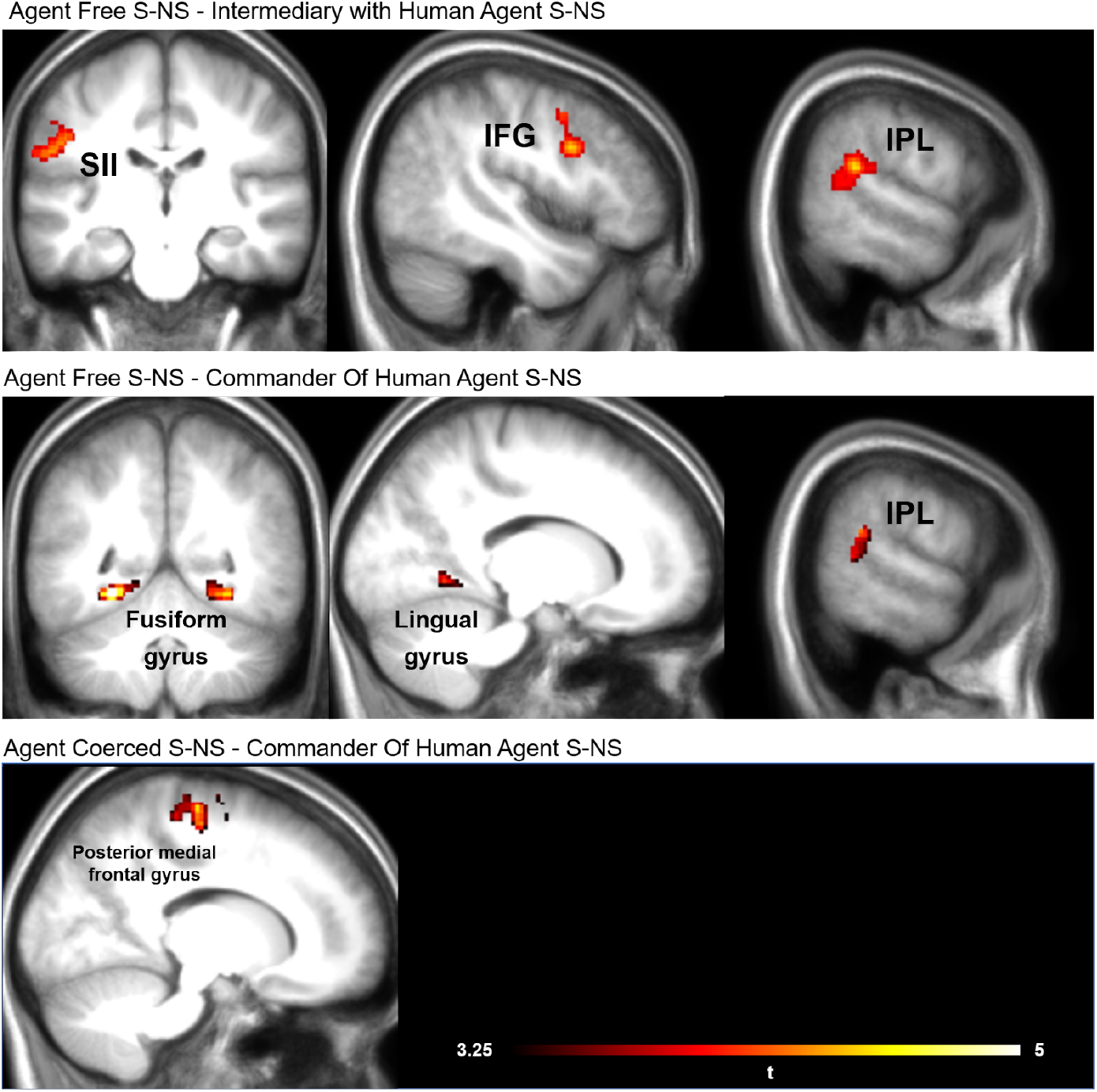
Results of two sample t-tests between the conditions [AgentFree(S-NS)-IntermediaryWithHumanAgent(S-NS)] _FWE_ at cluster level (236 voxels), t=3.5, *p* < .001,[AgentFree(S-NS)-CommanderOfHumanAgent(S-NS)] _FWE_ at cluster level (163 voxels), t=3.5, *p* < .001and [AgentCoerced(S-NS)-CommanderOfHumanAgent(S-NS)] _FWE_ at cluster level (315 voxels), t=3.5, *p* < .001. Peak coordinates can be seen in Table S2. None of the reverse contrasts demonstrated any significant results.

### Vicarious Pain Signatures

To examine if the lack of difference between our three conditions was due to the strict criteria of mass multivariate testing in fMRI, and to explore more specifically whether the manipulation influences empathic brain responses, we leverage the multivariate signature for vicarious pain developed by Zhou et al., (2020) to quantify empathic responses while seeing bodyparts in pain. We chose this particular signature, because it was trained on images of bodyparts in pain that best approximates the sight of hand receiving shocks in our study. **Fig. 5** shows the differential response (Shock-NoShock) of this signature in the conditions of the current study and in the study of Caspar, Ioumpa et al., 2020. As expected, in all cases, the differential response was significantly positive. At the second level, we thus conducted six two-sample t-tests comparing the two conditions of the previous agent study with the three conditions of the current commander study as follows: AgentFree(S-NS) - CommanderOfRobotAgent(S-NS) (t_(52)_=1.195, *p*=.237, Cohen’s d=.327, BF_10_=.496), AgentFree(S-NS) - CommanderOfHumanAgent(S-NS) (t_(52)_=1.546, *p*=.128, Cohen’s d=.423, BF_10_=.733), AgentFree(S-NS) - IntermediaryWithHumanAgent(S-NS) (t_(52)_=2.508, *p*=.015, Cohen’s d=.686, BF_10_=3.444), AgentCoerced(S-NS) - CommanderOfRobotAgent(S-NS) (t_(52)_=.108, *p*=.914, Cohen’s d=.30, BF_10_=.276), AgentCoerced(S-NS) - CommanderOfHumanAgent(S-NS) (t_(52)_=.587, *p*=.560, Cohen’s d=.160, BF_10_=.317), AgentCoerced(S-NS) - IntermediaryWithHumanAgent(S-NS) (t_(52)_=1.609, *p*=.114, Cohen’s d=.440, BF_10_=.795). These analyses revealed enhanced activation when agents were freely deciding compared to intermediates that were following orders and then delivering the same orders to a human agent. The other comparisons showed evidence for absence or close to evidence of absence of an effect. We additionally performed three paired-sample t-tests comparing the three conditions of the current commander study: IntermediaryWithHumanAgent(S-NS) - CommanderOfHumanAgent(S-NS) (t_(21)_=-.723, *p*=.477, Cohen’s d=-.151, BF_10_=.277), IntermediaryWithHumanAgent(S-NS) - CommanderOfRobotAgent(S-NS) (t_(21)_=-1.274, *p*=.216, Cohen’s d=-.266, BF_10_=.448), CommanderOfHumanAgent(S-NS) - CommanderOfRobotAgent(S-NS) (t_(21)_=-.447, *p*=.659, Cohen’s d=-.093, BF_10_=.240) which showed evidence for absence of a difference between the conditions.

**Fig. 5.**
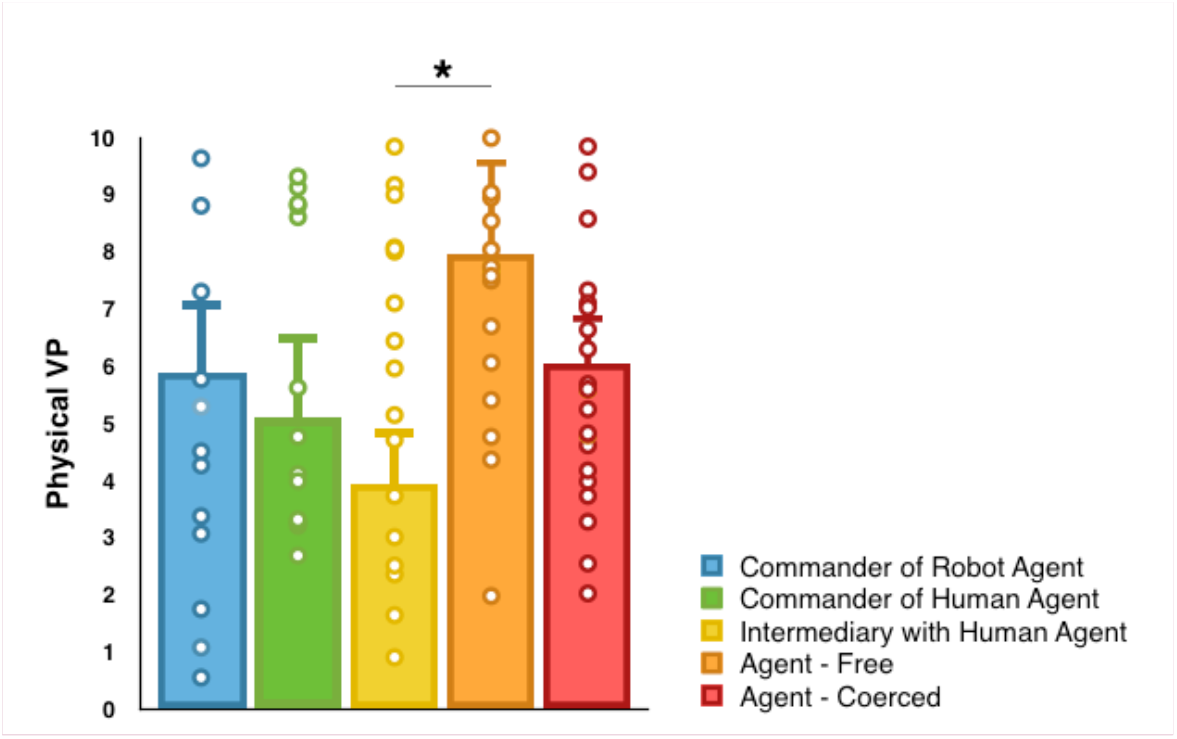
Results from the neurological physical vicarious pain signature analysis for the conditions Agent Free S-NS, Agent Coerced S-NS, IntermediaryWithHumanAgent S-NS, CommanderOfHumanAgent S-NS and CommanderOfRobotAgent S-NS. All comparisons were two-tailed.

## DISCUSSION STUDY 1

In Study 1, we aimed to understand if commanding or being in the position of the intermediary transmitting orders would influence how participants process the pain of a victim receiving mildly painful electric shocks. We also sought to understand how giving orders to another human being or to a robot would influence the same process. The BOLD signal clearly distinguished shock and no-shock trials in regions typically associated with empathy for pain, including cingulate, insular and somatosensory brain regions (Jauniaux et al., 2019; Lamm et al., 2011). Results from our *f*MRI analyses however did not reveal differences between our three conditions sufficiently strong to survive our p<0.001 threshold. With N=23 participants, and p<0.001 voxelwise threshold, our study would require such differences to have a large effect size of d=0.9 to be detected in 80% of cases so that our lack of significant difference suggests the absence of a large effect size of our manipulation. Smaller effect sizes of our manipulation cannot be excluded.

To explore the notion that all of the conditions tested here lead to reduced pain processing compared to directly being the agent, we ran additional comparisons between the remote conditions in the current experiment with previous data using a matching experimental design in which participants were the agent delivering the shocks (Caspar, Ioumpa et al., 2020). Specifically, when comparing the conditions in which participants were intermediaries (i.e. IntermediaryWithHumanAgent condition) to the condition in which participants were agents and free to decide, we observed higher activation when witnessing Shock vs NoShock outcomes for the Free condition in SII, IFG and IPL, which are key brain regions for empathy and social cognition (Shamay-Tsoory, 2011; (Janowski et al., 2013; Lockwood et al., 2013; Schulte-Rüther et al., 2007; Zaki, 2014). Additional evidence for different processing between these two conditions also came from our neurological signature analysis. We then compared the conditions in which participants could freely decide which orders to give to another human (i.e. CommanderOfHumanAgent) to the equivalent free situation, in which participants were agents and free to decide which button to press (i.e., Free condition, Caspar, Ioumpa et al., 2020). Results indicated that activity in areas IPL and fusiform gyrus, that have been linked in the literature with empathy and emotional social perception (Geday et al., 2003; Zaki et al., 2009; Janowski et al., 2012) was higher when participants were agents and could freely decide than when they were commanders and could freely decide. Comparing free agents and free commanders was interesting as the decisional power is the same for both roles when they are free to decide about the action to perform, but only agents execute the motor actions leading to the outcome. The differences in activation observed between free agents and free commanders thus suggest that performing the action engages more areas that are important for social cognition compared to having decisional power but being further away from the outcome of that same action.

Comparing the CommanderOfHumanAgent condition and the Agent Coerced (from Caspar, Ioumpa et al., 2020) showed more activation for agents than for commanders in posterior medial frontal gyrus, an area that has been linked with cognitive control, response conflict, decision uncertainty and cognitive dissonance (Ridderinkhof et al., 2004, Izuma et al., 2015). This suggests that in hierarchical situations, the agent is more engaged and experiencing more conflict for his actions, even coerced ones, than commanders giving orders. However it is important to be aware of the limitations of reverse inference and the difficulty of unambiguously associating activity in specific brain regions with mental processes (Poldrack, 2006). Thus these conclusions should be interpreted with caution.

All our participants acted both as agents and victims, in randomized order. To examine if having been a victim altered the behavior of commanders compared to being commander first, we examined whether the number of shocks and the sense of responsibility was influenced by the order. In either case we did not find evidence for an effect of order, and we thus did not further consider the effect of orders for the fMRI analysis. Importantly for the fMRI analyses, we also did not measure significant differences in the number of shocks delivered to the victims across our three commander conditions, which simplifies the interpretation of the fMRI data. The number of shocks given also did not differ significantly between the current experiment and the agent experiment reported in Caspar, Ioumpa et al., 2020 (CI_95_=-0.995-0.091, t_(53)_=-1.661, *p*=.103, Cohen’s d=-0.454, BF_10_=.851 for the HumanAgent comparison and CI_95_=-1.059, t_(53)_=-1.892, *p*=.064, Cohen’s d=-0.517, BF_10_=1.183 for the RobotAgent comparison).

## STUDY 2 – METHOD

### Participants

Forty-eight participants (24 males, 24 females) were recruited in 24 dyads. None of the participants reported to know each other. The mean age was 23.90 (SD=3.93). We recruited a larger sample than in Study 1 because we expected to have to reject more participants because the testing took place in a month of the year involving very hot temperatures and the EEG data were particularly difficult to acquire due to sweat artifacts. The following exclusion criteria were determined prior to further analysis: (1) failure to understand the task, (2) failure to perform correctly the task measuring the implicit sense of agency, (3) failure to obtain a good signal-to-noise ratio for electroencephalography (EEG) recordings or (4) failure to report the presence of pain on the victim’s hand with the pain scale. No participants were excluded for this reason. To identify participants for whom the estimated action-tone intervals did not gradually increase with the real action-tone intervals, we performed Pearson correlations. When the Pearson r was lower than .1, we excluded the action-tone intervals for the corresponding participant. We also did not analyze the action-tone intervals data of participants who sent less than 5/60 shocks or more than 55/60 shocks in at least one of the experimental conditions since it would lead to unreliable statistical comparison between Shock and No shock trials. Accordingly, the action-tone intervals of 7/48 participants were lost due to a Pearson r < .1. 14/48 participants sent less than 5/60 shocks or more than 55/60 shocks in at least one of the experimental conditions and their action-tone interval data were lost. The action-tone interval data of one participant was included in the two categories (i.e., r < .1 and unreliable number of shock/no shock trials). As a result, we lost the action-tone interval data of 20/48 participants. The EEG data of 18 participants were not analyzed: 4 because of too many visual artifacts, head artifacts and/or sweat artifacts, and 14 because they delivered only a small number of shock (<5/60, N=10) or a high number of shocks (>55, N=4) in either one or all conditions. This would indeed prevent obtaining a reliable difference between shock and no shock trials (Caspar, Ioumpa, et al., 2020). The study was approved by the local ethics committee of the local ethical committee of the Université libre de Bruxelles (ref: 018/2015). Data are made available on OSF (*will be made public after publication*).

### Method and Material

The method was globally similar to Study 1, including the same conditions (Fig. 1A) but with a slightly different timing of trials. Each trial started with a fixation cross lasting between 1 and 2 seconds. When the fixation cross disappeared, participants received a verbal instruction from the experimenter, similarly to Study 1. Then, they had to press one out of two buttons: SHOCK or NO SHOCK in order to send an order to an agent, either human or robot. After the agent pressed the button corresponding to the order of the participant, a tone was presented (400Hz, 200ms). The interval between the agent’s keypress and the start of the beep was of 200, 500 or 800 ms. Participants were asked to estimate the elapsed time between their own keypress when they sent the order and the beep onset. If a shock was sent to the ‘victim’, the shock was delivered at the exact same time as the tone to avoid temporal bias. After 2 seconds, an analogue scale with ‘0’ on the left side and ‘1,500’ on the right side of the scale was then displayed on the screen. A red position marker was displayed on that scale with a number, corresponding to the marker’s current position in *ms*. The starting position of the marker varied randomly on a trial-wise basis and participants were told to ignore the starting position of the marker to provide their final answer. Participants could move the position of the marker along the analogue scale by using the same two buttons as for shock and no shock. The keys below the middle fingers allowed to modify the number associated with the position of the rectangle by steps of +-100 ms. The keys below the index fingers allowed to modify the answer by steps of +-1 ms. After a fixed duration of 6 seconds, their answer was saved and the next trial started. Each participant started with a training session to practice the time interval procedure. The training session lasted for minimum 8 trials and was repeated until participants declared that they could perform the task correctly.

To preserve the same experimental set-up between Study 1 and Study 2, participants were isolated in a room and ‘victims’ were in another room with the camera displaying their hand in real-time on the participants’s screen. In all three experimental conditions, the experimenter came to talk to the participant before the start of each experimental condition but then left the room by mentioning that it was to avoid too many interferences in the EEG recordings due to her presence. participants were told that they would hear the experimenter’s instructions through the headphones.

Each experimental condition was composed of 60 trials. Order of the experimental conditions was counterbalanced across participants. The same questionnaires were presented to participants at the end of the experimental session.

### EEG recordings

Brain activity was recorded using a 64-channels electrode cap with the ActiveTwo system (BioSemi) and data were analyzed using Fieldtrip software (Oostenveld et al., 2011). The activities from left and right mastoids and from horizontal and vertical eye movements were also recorded. Amplified voltages were sampled at 2048 Hz. Data were referenced to the average signal of the mastoids and filtered (low-pass at 50 Hz and high-pass at 0.01 Hz). Artifacts due to eye movements were removed based on a visual inspection with the removal of epochs containing eye blinks or ocular saccades. Because of the EEG recordings, participants were further instructed to wait a minimum of 1 s in a relaxed position before pressing a key, so as to obtain a consistent and noise-free baseline taken -500 to -300 ms before the occurrence of the tone. Participants were additionally instructed not to move for up to 2 s after the tone and asked to avoid blinking when they pressed a button. To ensure that participants respected the 2 s without moving and blinking after the tone, they were told to wait for the time scale to appear on the screen. Trials in which participants disobeyed the orders of the experimenter were removed from the analysis.

All event-related potentials were analyzed across Fz, FCz, Cz, CPz, and Pz. The N1 and the N2 were measured as the most negative peaks within the 30-130ms time-window and the 240-340 ms time-window after the tone, respectively. The P2 and the P3 were measured as the most positive peaks within the 130-230 ms time-window and the 340-440ms time-window after the tone, respectively. The early LPP and the late LPP were measured as the mean amplitude between the 440-650 ms time-window and the 650-900 ms time-window after the tone, respectively.

Source reconstruction was conducted on the Grand Average of EEG data that were computed with a Noise Covariance Estimation. Minimum norm estimation (MNE; Dale & Sereno, 1993) was applied to reconstruct the sources of ERPs components. The volume construction was based on standard head model and source model downloaded through Fieldtrip. Having performed the source localization, we used it to create brain maps showing the brain regions involved in the activity associated with each ERP. We performed this operation with a custom made python software that uses as input the results of our source localization (https://github.com/ldeangelisphys/ft2nii/). The software assigns each localized source to the corresponding voxel in a standard 2 mm MNI template (MNI152), performs a temporal average over the time window corresponding to the ERP of interest, and a spatial smoothing of the resulting map with a 5 mm gaussian kernel.

## RESULTS – STUDY 2

### Number of shocks delivered

In the IntermediaryWithHumanAgent, participants were ordered by the experimenter, on a trial-basis, to tell the human agent to inflict 30/60 shocks to the ‘victim’, randomly. In the Commander conditions, participants could freely decide which order to send to the agent. Descriptive statistics indicated that in the CommanderOfHumanAgent condition, participants asked the agent to administer 22.46/60 (SD=17.80, min: 0 – max: 60) shocks to the ‘victim’. In the CommanderOfRobotAgent condition, participants asked the agent to administer 22.56/60 (SD=17.90, min: 0 – max: 60) shocks to the ‘victim’. In the IntermediaryWithHumanAgent, participants transmit the order to send a shock to the ‘victim’ on 27.54/60 trials (SD=7.59, min: 0 – max: 33). Thirteen out of 48 participants reported that they voluntarily disobeyed the orders of the experimenter on some trials in the IntermediaryWithHumanAgent condition. Among those 13 participants, 10 disobeyed prosocially, that is, by refusing to send a shock during ‘shock trials’ and by telling the agent not to deliver a shock to the ‘victim’ even if the experimenter asked them to do so (i.e. prosocial disobedience), and 3 disobeyed ‘by contradiction’, that is, they disobeyed as often on ‘don’t shock’ trials and on ‘shock trials’. We conducted a repeated-measures ANOVA with Condition (CommanderOfHumanAgent, CommanderOfRobotAgent, IntermediaryWithHumanAgent) as within subject factor and Role-Order (Commander first, Victim first) as between-subject factor on the number of shocks sent in each experimental condition. We observed a significant main effect of Condition (F_(2,92)_=4.119, *p*=.019, η^2^_***partial***_ = .082, BF_incl_=1.472), see **Fig. 6A**. Paired-comparisons indicated that participants administered less shocks in the CommanderOfHumanAgent condition and in the CommanderOfRobotAgent condition than in the IntermediaryWithHumanAgent condition (t_(47)_=2.130, *p*=.038, Cohen’s d=.307 and t_(47)_=2.051, *p*=.046, Cohen’s d=.296, respectively). Both the frequentist and the Bayesian results indicated that Order of the role was slightly in favor of H_0_ (*p*>.3, BF_incl_=.391). The interaction Condition * Order of the role was in favor of H_0_ (*p*>.2, BF_incl_=.280).

**Fig 6.**
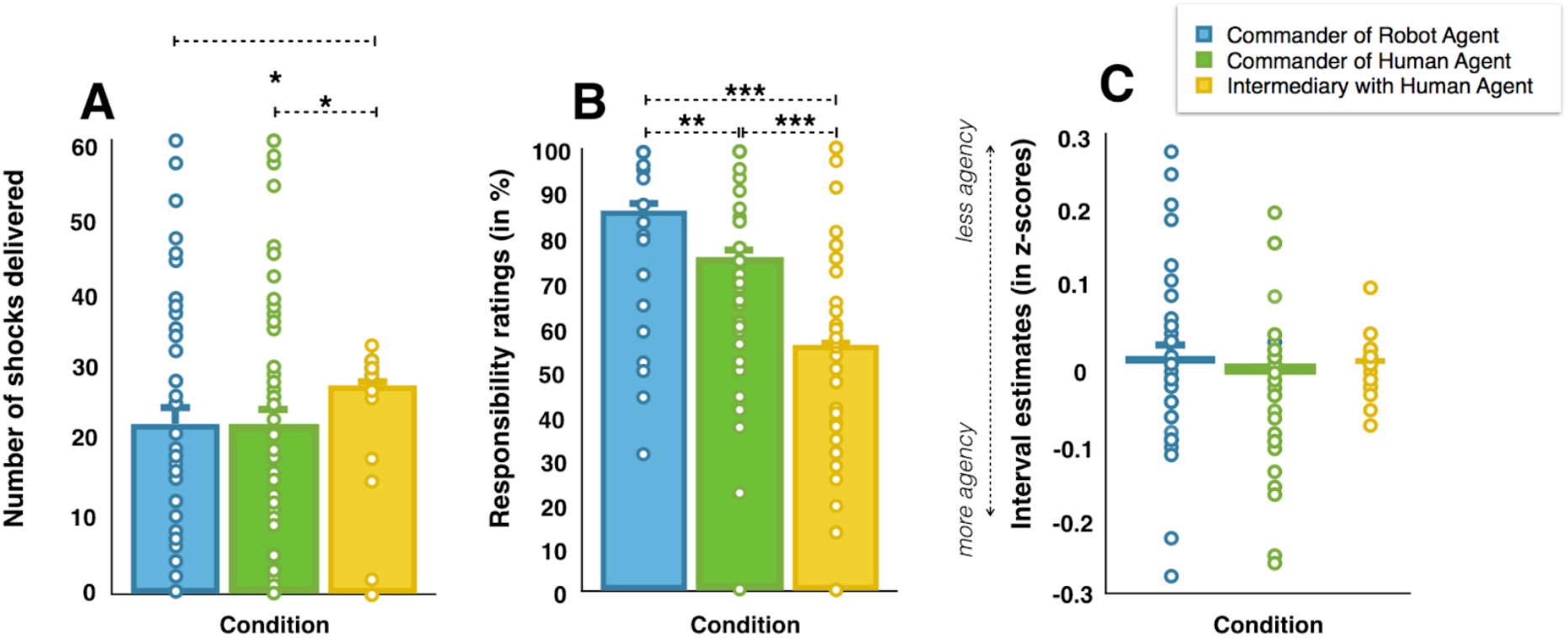
A) Graphical representation of the number of shocks delivered in the three experimental conditions. B) Graphical representation of responsibility ratings in the three experimental conditions. C) Graphical representation of z-scores of interval estimates in the three experimental conditions. All tests were two-tailed. *** represents p <=.001 and a BF > 3.

### Pain Scale

The estimations reported by agents on the pain scale were not analyzed as only 9/23 participants had data for shock trials in the three experimental conditions.

### Responsibility ratings

At the end of each experimental condition, participants had to report how responsible they felt for the outcome of their orders. We conducted a repeated-measures ANOVA with Condition (CommanderOfHumanAgent, CommanderOfRobotAgent, IntermediaryWithHumanAgent) as within subject factor and Role-Order (Commander first, Victim first) as between-subject factor on the responsibility ratings. Both the frequentist and the Bayesian results supported a main effect of Condition (F_(2,92)_=28.917, *p* < .001, η^2^_***partial***_ = .386, BF_incl_=1.234E+8), see **Fig. 6B**. Paired-comparisons indicated that responsibility ratings were higher in the CommanderOfRobotAgent condition (86.2%, CI_95_=80.9-91.5) than in the CommanderOfHumanAgent condition (75.6%, CI_95_=68.7-82.5, t_(47)_=3.280, *p* = .002, Cohen’s d=.473, BF_10_=15.87) and, than in the IntermediaryWithHumanAgent condition (55.8%, CI_95_=47.6-54, t_(47)_=-6.985, *p* < .001, Cohen’s d=-1.008, BF_10_=1.456E+6). Responsibility ratings were also higher in the CommanderOfHumanAgent condition than in the IntermediaryWithHumanAgent condition (t_(47)_=-4.538, *p* < .001, Cohen’s d=-.655, BF_10_=545). The order of the role (F_(1,46)_=.140, *p* > .7, BF_incl_=.208) and the interaction (F_(22,92)_=.077, *p* > .9, BF_incl_=.106) were in favor of H_0_ and thus did not show evidence for influencing responsibility ratings.

### Sense of agency

Because interval estimates were planned to be correlated with other measurements in future analyses, we first transformed the raw interval estimates in z-score data. It is indeed known that participants may differ in the way they use the ms-scale to provide an answer, some preferring smaller numbers and others preferring larger numbers (Caspar, Lo Bue, et al., 2020; Cravo et al., 2013). Z-scores reduce irrelevant inter-subject variability by subtracting from each interval estimate, the mean estimate for that participant across all trials and by dividing the resulting differences by the standard deviation of all estimates for that participant. The z-scored interval estimates are interpreted as the raw interval estimates are, with lower z-scored interval estimates being interpreted as a higher sense of agency (SoA). Trials where participants disobeyed the orders from the experimenter were removed from the analyses. We conducted a repeated-measures ANOVA with Condition (CommanderOfHumanAgent, CommanderOfRobotAgent, IntermediaryWithHumanAgent) and Shocks (Shock, No shock) as within subject factors and Role-Order (Commander first, Victim first) as between-subject factor on z-scores. The main effect of Condition and Order of the role were in favor of H_0_, respectively *p* > .3, BF_incl_=.036 and *p* > .9, BF_incl_=.115, see **Fig. 6C**. The main effect of Shock was inconclusive (*p* > .1, BF_incl_=1.547). All the interactions were in favor of H_0_ (all *p*s > .1, all BFs_incl_ ≤ .199). We further ran exploratory analyses in order to investigate if self-reported personality traits influenced z-scored interval estimated when commanding freely or being an intermediary. We thus computed a “Commander effect”, which is the difference between the IntermediaryWithHumanAgent condition and the corresponding CommanderOfHumanAgent conditions (i.e. IntermediaryWithHumanAgent - CommanderOfHumanAgent). With this subtraction, higher positive values indicated more SoA in the CommanderOfHumanAgent condition than in the IntermediaryWithHumanAgent condition. We observed a small evidence that the less participants scored on the ASC submission scale, the lower their interval estimates were in the CommanderOfHumanAgent condition compared to the IntermediaryWithHumanAgent condition (r=-.392, *p* = .022, BF_10_=2.628). Correlations with the other subscale were inconclusive or in favor of H0 (all ps > .042, all BFs10<1.511).

### EEG results

We compared the neural processing of pain with an electroencephalogram when participants witnessed a shock being delivered on the hand of the ‘victim’. We extracted, based on both previous literature and the visual inspection of the grand averaged waves, the amplitude of several event-related potentials associated either with auditory outcome processing (N1, P2, Luck, 2012) or with pain outcome processing (N2, P3, eLPP, lLPP, Chen et al., 2014), see **Fig. 7A**.

**Fig 7.**
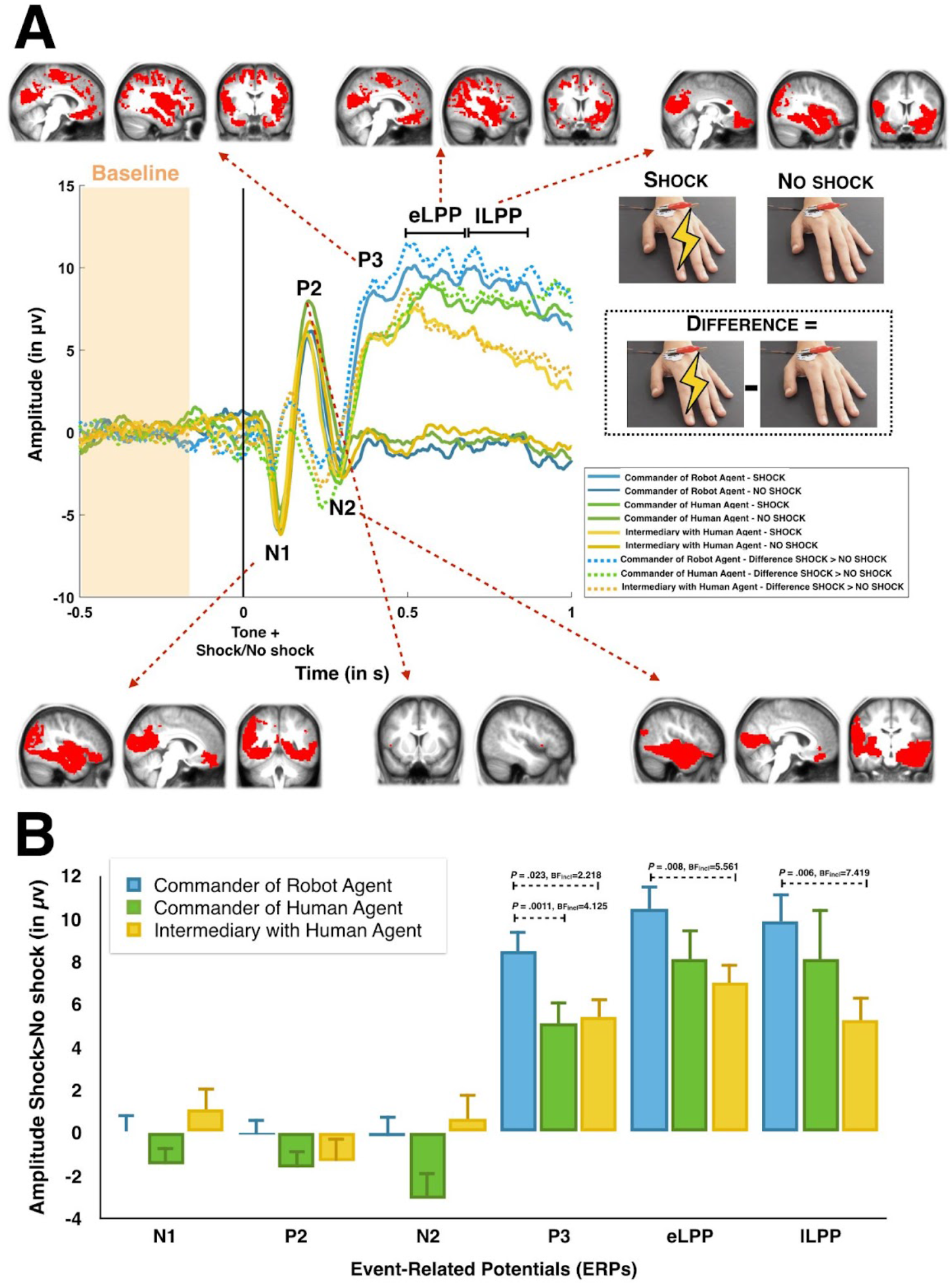
A) Grand average ERP for the Shock trials (light full lines) and the NoShock trials (dark full lines). The dotted lines represent the difference Shock - NoShock trials. Source reconstruction maps for each ERP are displayed along each ERP for the difference Shock - No trials irrespectively of condition for the voxels with the highest 5% of positive values. B) Graphical representation of the amplitude in μV of each ERP in each experimental condition. All tests were two-tailed. Only significant pairwise comparisons are shown, see Table 2 and text for the other comparisons). Source reconstruction maps represent the highest 5% of activation.

We first compared the amplitude of those potentials when participants witnessed a shock on the victim’s hand to when they did not witness that shock, in order to identify the ERPs sensitive to seeing pain. To do so, we averaged the amplitude of each ERP across the three experimental conditions for shock and no shock trials. Results supported evidence for a higher amplitude for shock trials in comparison with no shock trials for the P3 (t_(30)_=10.108, *p* < .001, Cohen’s d=1.815, BF_10_=2.673E+9), the early LPP (t_(30)_=10.890, *p* < .001, Cohen’s d=1.956, BF_10_=1.443E+9) and the lLPP (t_(30)_=7.044, *p* < .001, Cohen’s d=1.265, BF_10_=187382.55). This difference was in favor of H_0_ for the N1 (*p* > .8, BF_10_=.195) and for the N2 (*p* > .3, BF_10_=.308) and inconclusive for the P2 (*p* =.048, BF_10_=1.214). Those results confirmed that the P3, the early LPP and the late LPP were sensitive to the visualization of a painful stimulus delivered on the hand of the victim.

In order to evaluate how the experimental conditions influenced the neural response to the pain of the ‘victim’, we then performed a repeated-measures ANOVA with Condition (CommanderOfHumanAgent, CommanderOfRobotAgent, IntermediaryWithHumanAgent) as a within-subject factor and Role-Order (Commander first, Victim first) as a between-subject factor on the computed difference between shocks and no shocks trials for those potentials showing a Shock-NoShock effect (i.e. P3, early LPP and the late LPP). For the P3, we observed a main effect of condition (F_(2,58)_=4.502, *p* = .015, η^2^_***partial***_ = .130, BF_incl_=3.0958), see **Fig. 7B**. Paired comparisons supported that the amplitude of the P3 was higher in the CommanderOfRobotAgent condition than in the IntermediaryWithHumanAgent condition (t_(30)_=-2.712, *p* = .0011, Cohen’s d=-0.487). **Table 2** displays the results of the paired comparisons between conditions. We found evidence in favour of H_0_ for the main effect of Order of the role (*p* > .7, BF_incl_=.251) and for the interaction (*p* > .3, BF_incl_=.264). For the early LPP, the main effect of Condition was significant (F_(2,58)_=3.465, *p* = .038, η^2^_***partial***_ = .099) but inconclusive with the Bayesian approach (BF_incl_=1.197). We found evidence in favour of H_0_ for the main effect of Order of the role (*p* > .9, BF_incl_=.318) and slightly in favour of H_0_ for the interaction (*p* = .09, BF_incl_=.564). For the late LPP, the main effects of Condition and Order of the role were inconclusive (*p* > .09, BF_incl_=0.709 and *p* > .5, BF_incl_=0.428, respectively). The interaction was significant with the frequentist approach (F_(2,58)_=3.596, *p* = .034, η^2^_***partial***_ = .102) but inconclusive with the Bayesian approach (BF_incl_=0.977).

**Table 2.** Pairwise comparisons of Shock-NoShock difference across conditions. For the three ERPs that are sensitive to shocks, the table summarizes the pairwise comparison across conditions. Means detail the mean voltage (in microvolts) per condition, SDs their standard deviation across participants, Df, t and p summarize the two-tailed student t-test, BF10 the Bayesian equivalent. Intermed = IntermediaryWithHumanAgent condition; CH = CommanderOfHumanAgent condition; CR = CommanderOfRobotAgent condition. Green fonts indicate results in favour of H1 with both the frequentist and the Bayesian approaches. Red fonts indicate results in favour of H0 (non-significant frequentist statistics and BF10<⅓).

| ERP | Comparisons | Means | SDs | df | t | p | Cohen's d | BF<sub>10</sub> |
| --- | --- | --- | --- | --- | --- | --- | --- | --- |
| P3 | Intermed - CH | 5.4μv - 5.1μv | 4.7 - 5.9 | 30 | 0.278 | 0.783 | 0.050 | 0.199 |
|  | Intermed - CR | 5.4 - 8.5 | 4.7 - 5.2 | 30 | -2.396 | 0.023 | -0.430 | 2.218 |
|  | CH - CR | 5.1 - 8.5 | 5.9 - 5.2 | 30 | -2.712 | 0.011 | -0.487 | 4.125 |
| eLPP | Intermed - CH | 7 - 8.1 | 5.1 - 7.7 | 30 | -0.654 | 0.518 | -0.117 | 0.233 |
|  | Intermed - CR | 7 - 10.5 | 5.1 - 5.8 | 30 | -2.857 | 0.008 | -0.513 | 5.561 |
|  | CH - CR | 8.1 - 10.5 | 7.7 - 5.8 | 30 | -1.888 | 0.069 | -0.339 | 0.920 |
| ILPP | Intermed - CH | 5.2 - 8.1 | 6.3 - 13 | 30 | -1.059 | 0.298 | -0.190 | 0.320 |
|  | Intermed - CR | 5.2 - 9.8 | 6.3 - 7.3 | 30 | -2.992 | 0.006 | -0.537 | 7.419 |
|  | CH - CR | 8.1 - 9.8 | 13 - 7.3 | 30 | -0.790 | 0.436 | -0.142 | 0.255 |

#### Source reconstruction results

We computed the maps corresponding to our three contrasts of interest[CommanderOfHumanAgent(S-NS)-IntermediaryWithHumanAgent(S-NS)],[CommanderOfRobotAgent(S-NS)-IntermediaryWithHumanAgent(S-NS)],[CommanderOfRobotAgent(S-NS)-CommanderOfHumanAgent(S-NS)]. We then post-processed them by selecting the voxels with the highest 5% of values encoding for regions that are the most positively involved in the selected ERP. Results shown in **Fig. 8**.

**Fig. 8:**
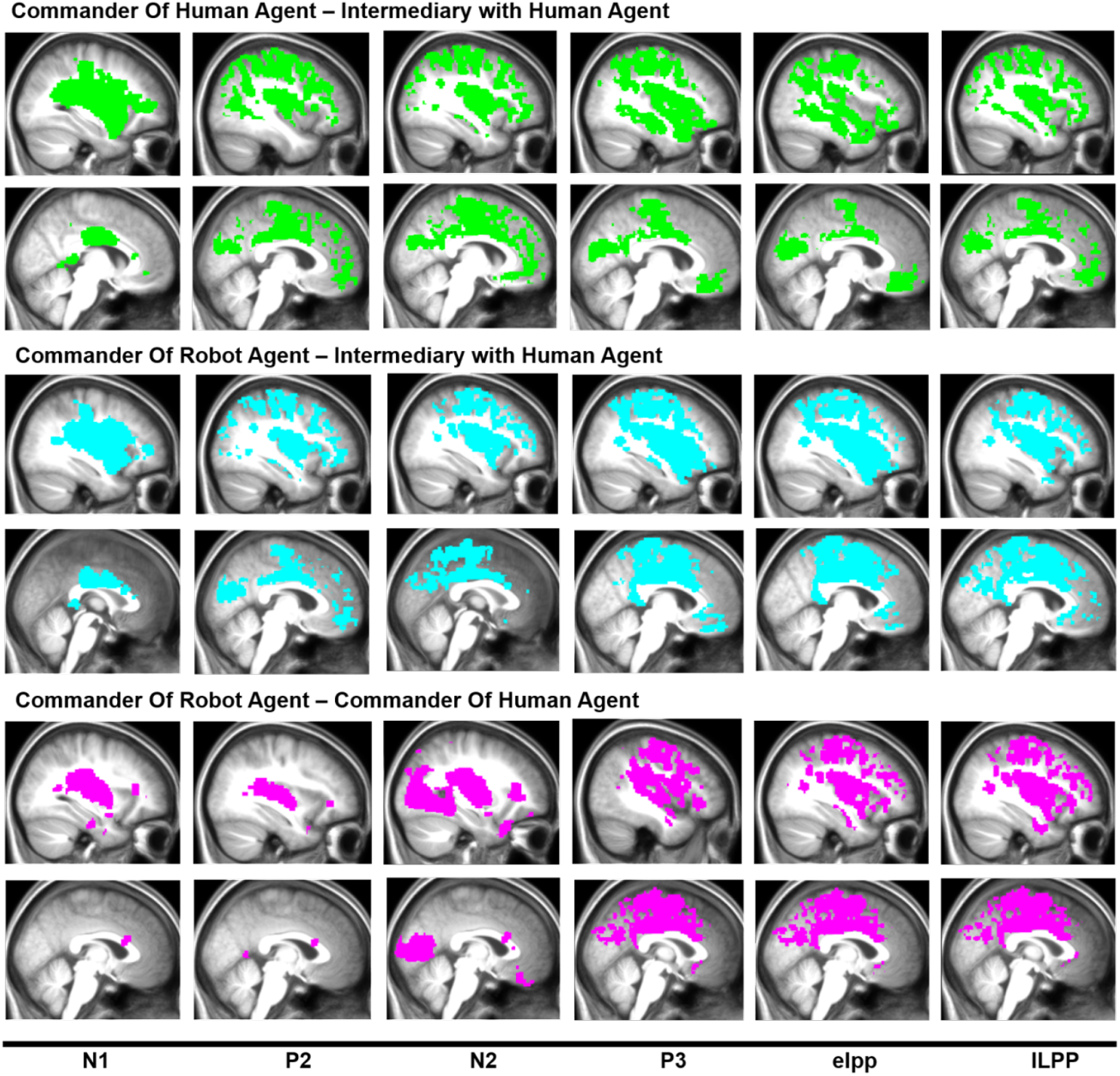
maps derived reconstructing the EGG signal for the contrasts [CommanderOfHumanAgent(S-NS)-IntermediaryWithHumanAgent(S-NS)] in green, [CommanderOfRobotAgent(S-NS)-IntermediaryWithHumanAgent(S-NS)] in cyan, [CommanderOfRobotAgent(S-NS)-CommanderOfHumanAgent(S-NS)] in violet. The voxels with the highest 5% of values are displayed.

In order to compare Study 1 and Study 2 and to ensure that witnessing a shock versus no shock being delivered on the victim’s hand involve a similar brain activation pattern, we overlaid the MRI results obtained in Study 1 to the EEG results obtained in Study 2 for the Shock - No Shock contrast. Results are displayed in **Fig. 9**.

**Fig. 9:**
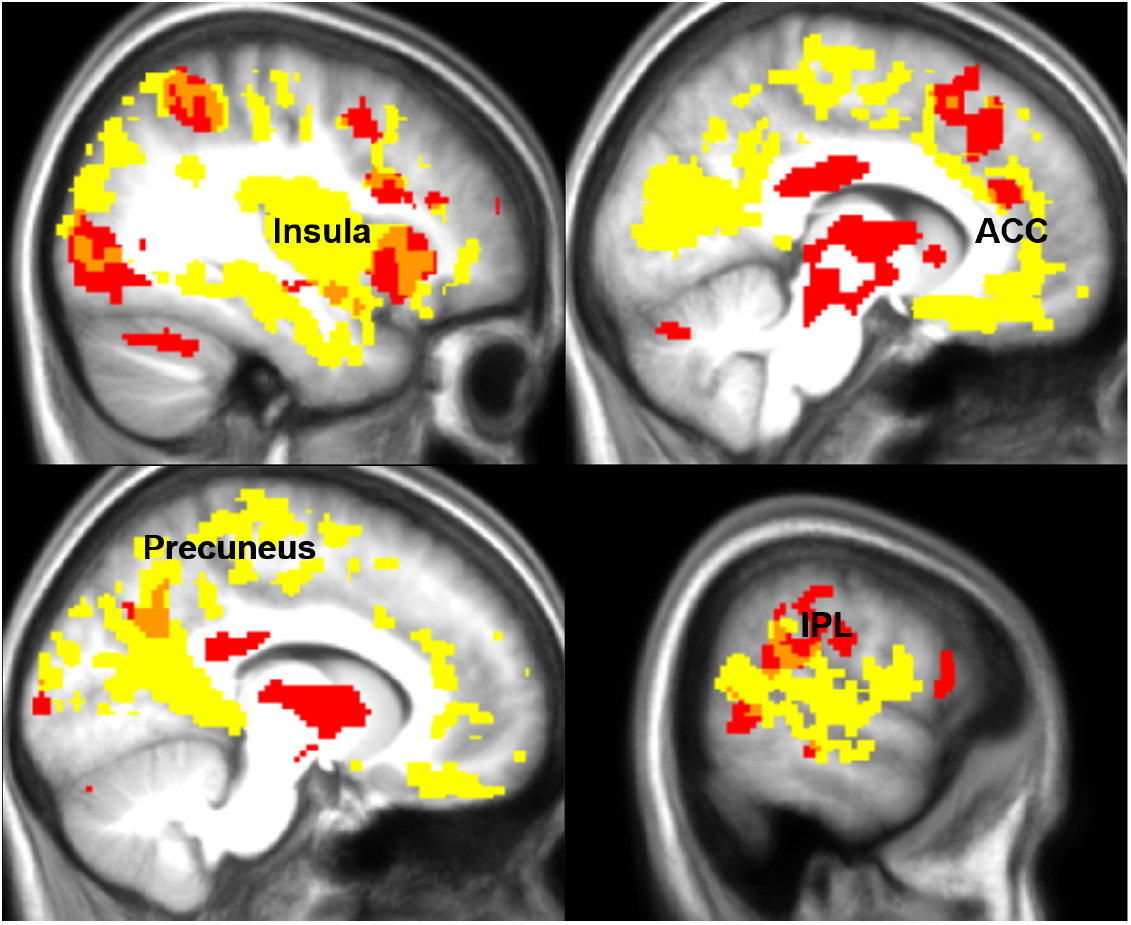
Overlap (in orange) between the maps from the GLM analyses for Sock-NoShock contrast (in red) and the 5% higher activation map derived reconstructing the EGG signal for the Sock-NoShock contrasts using the mean activation from components P3, eLPP and lLPP (in yellow).

### Relationship between number of shocks freely ordered and temporal binding, feeling of responsibility and ERP

In order to investigate to what extent the sense of agency, empathy for pain and feeling of responsibility drive prosocial behaviors, we further performed Pearson correlations. In order to create a single variable ‘free-choice condition’, irrespective of the type of agent, we summed the data of the CommanderOfHumanAgent and CommanderOfRobotAgent conditions for the number of shocks delivered. We then computed an average score across the same conditions for the z-scores of interval estimates, used as a proxy for the sense of agency, for responsibility ratings and for ERPs that were sensitive to the visualization of the victim’s pain (i.e. P3, eLPP and lLPP). For these ERPs, we computed a general pain response by subtracting the amplitude of those potentials during No shock trials to Shock trials (i.e. Shock-No shock). To correct for multiple comparisons with the frequentist statistics, we applied a False Discovery Rate (FDR) approach with the Benjamini and Hochberg method (Benjamin & Hochberg, 1995) to each p-value. Both frequentist and Bayesian statistics for those correlations were two-tailed. We observed evidence that the number of shocks freely administered to the victim correlated positively with the z-scores of interval estimates (r=.443, *p*_*FDR*_ = .022, BF_10_=3.311), indicating that the higher the z-scores were, which corresponds to a reduced sense of agency, the higher the number of shocks sent to the victim was, see **Fig. 10**. We also observed evidence for a negative correlation between the number of shocks freely delivered and responsibility ratings (r=-.434, *p*_*FDR*_ = .015, BF_10_=7.372). This suggests that the more responsible participants felt, the less shocks they sent to the victim. Correlations with ERPs revealed evidence for a negative correlation of the number of shocks ordered with the late LPP (r=-.561, *p*_*FDR*_ = .005, BF_10_=38.575) and the early LPP Shock-NoShock magnitudes (r=-.434, *p*_*FDR*_ = .022, BF_10_=3.857), suggesting that the higher the Shock-NoShock amplitudes of the early and late LPP were, the lower the number of shocks delivered. Of note, these results stayed similar when considering only the first half or the second half of the trials, thus controlling for the repetition suppression effect. The Bayesian approach indicated that the correlation between the number of shocks and P3 was slightly in favor of H0 (*p*_*FDR*_>.2, BF_10_=.384). Of note, the number of shocks delivered freely to the victim did not correlate with the amplitude of ERPs which were not found to be sensitive to the pain of the victim (i.e. N1, P2, N2 – all *p*s_*FDR*_>.072, BF_10_≥.609 & ≤1.924).

**Fig. 10.**
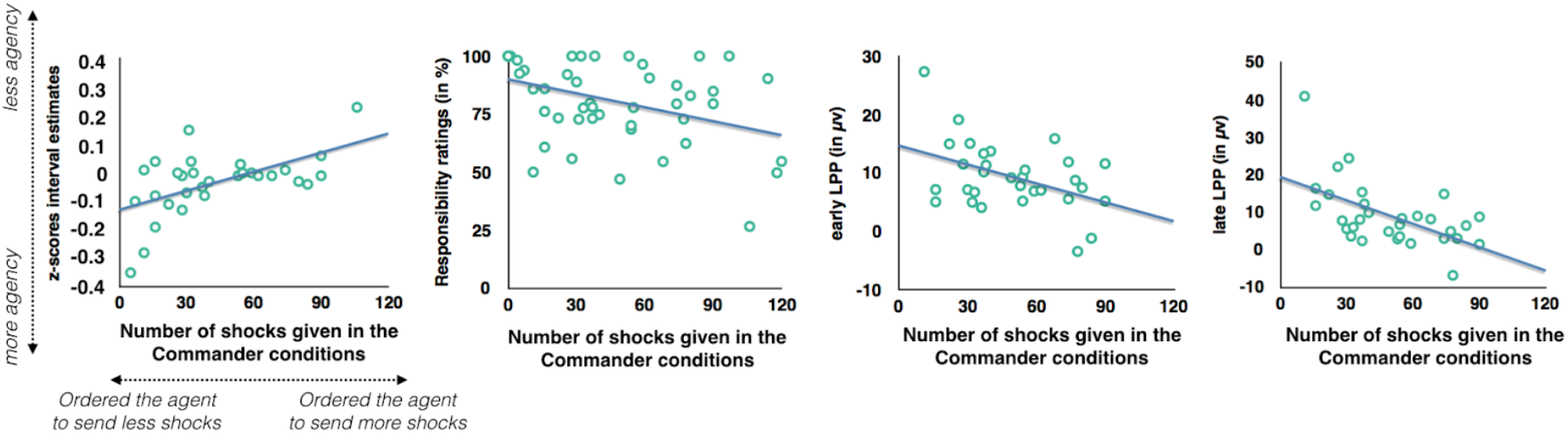
Graphical representations of Pearson correlations between the number of shocks given in the commander conditions and the sense of agency (top left), responsibility ratings (middle left), amplitude of the eLPP (middle right) and amplitude of the lLPP (top right). All tests were two-tailed.

## STUDIES 1 & 2 – Self-report personality Questionnaires

We conducted correlational, exploratory analyses on both studies combined in order to investigate self-reported personality traits associated with the number of shocks freely delivered in the Commander conditions, irrespective of the types of agent. Since the analyses were exploratory, analyses were two-tailed. Again, we corrected for multiple comparisons with the False Discovery Rate (FDR). Results revealed that the higher participants scored on the Levenson Primary Psychopathy scale, the more shocks they freely ordered the agent to send to the victim (r=.370, *p*_*FDR*_=.001, BF_10_=61.170). We also observed evidence for two positive correlations between the number of shocks delivered and the purity (r=.337, *p*_*FDR*_=.011, BF_10_=20.067) and the authoritarianism (r=.304, *p*_*FDR*_=.027, BF_10_=7.550) subscales of the Moral Foundation Questionnaires. Other correlations were in favor of H0 or inconclusive (all BFs_10_ > .137 & <1.61).

## DISCUSSION STUDY 2

We did not observe statistical differences in interval estimates, used as a proxy for the sense of agency, between the three experimental conditions. This suggests that the sense of agency (unlike the sense of responsibility) does not differ between commanding and being a mere intermediary. However, with the present results we cannot argue in favor of an equally low sense of agency when commanding or being a mere intermediary or in favor or a high sense of agency for both positions in the command chain. A control condition, in which participants are the direct agent of the action would have allowed to understand whether or not the sense of agency is reduced when people give orders to a third party. Yet, in a former study in which participants took either the role of the agent or the commander in a within-subject design (Caspar et al., 2018), results indicated that commanding an agent lead to a reduction of the sense of agency, also measures with the method of interval estimates. We further observed that self-reported personality traits modulated this effect, with participants scoring higher on the ASC scale having the lower commander effect.

While several scientific publications highlighted the role of the sense of agency in prosocial attitudes (Gallagher et al., 2002; Haggard, 2017), direct correlations between interval estimates and prosocial behaviors had barely been shown. Here, we observed a correlation between interval estimates and prosocial behaviors in the Commander conditions, irrespective of the types of agent (i.e. robot or human). This suggests that participants with a high sense of agency when sending orders to an agent tend also to act more prosocially, by avoiding to inflict pain to the ‘victim’ too frequently. Results further showed that participants which experienced a higher feeling of responsibility when commanding an agent also sent less shocks to the victim. This highlights the role of experiencing oneself as the author of an action and feeling responsible for its outcomes in prosocial decision-making.

In accordance with former studies (see Coll, 2018 for a meta-analysis), we observed that, especially the P3, but also the early and late LPPs, were sensitive to the observation of the painful shock delivered to the victim’s hand. Regarding our experimental manipulations, we observed that participants had a higher neural response when they could command a robot compared to when they were intermediary. This result was observed on the P3, but not on the eLPP and the lLPP, which is consistent with another former study manipulating social power (Galang et al., 2021). The literature on the P3 and the LPP does not offer a concrete conceptual distinction between these two components, some authors arguing that the LPP is a simple extension of the P3 (e.g. Olofsson et al., 2008). However, taken together, these results suggest that the P3 and the LPP could be influenced by different social factors.

The N1 and N2 amplitudes did not statistically differ between painful and non-painful trials. A possibility is that the N1 and the N2 components also partially reflect the neural processing of auditory outcomes (i.e. tones) on centro-parietal sites (Luck & Kappenman, 2011), for which we did not expect difference for pain and non-painful stimuli. However, as Coll et al. (2018) indicated in a recent meta-analysis, even former studies which indicated a difference between the amplitude of the N2 between painful and non-painful stimuli, N1 and N2 are not reliably associated with vicarious pain observation.

Interestingly, while the P3 appeared to be more influenced than the LPP by social power, the LPP correlated with prosocial behaviors while the P3 did not. We indeed observed that the higher the amplitude of the early and late LPPs was, the fewer shocks participants ordered to send to the victim. This is in line with former studies which showed that a higher neural empathic response leads to more prosocial attitudes towards others (e.g. Gallo et al., 2018).

### GENERAL DISCUSSION

Past scientific research has shown that being an intermediary in a command chain was associated with a higher prevalence to accept immoral orders (Milgram, 1974). In the present study, our aim was to understand how two different neuro-cognitive processes, that are, the sense of agency and empathy for pain, differ between being the commander or a simple intermediary.

In a former fMRI study (Caspar, Ioumpa et al., 2020), we observed that for the agent directly delivering a shock, obeying orders reduced vicarious activations towards a victim’s pain compared to acting freely, suggesting that a reduced decisional power negatively impacted the neural empathic response. This result was also confirmed by another study (Galang et al., 2021), which showed that recalling a low social power situation did not lead to difference in the neural empathic response between painful and non-painful pictures while this difference was significant in a high social power condition. In the present study, when participants were in the role of commander, they had a total social power as they could decide which order to ask an intermediary to execute. In contrast, when they were in the intermediary position, they had to follow the experimenter’s instructions, thus having a reduced social power.

Interestingly, in Study 1 we observed that vicarious activations towards the victim’s pain, as measured using a vicarious pain signature, or less directly, using voxelwise differences in regions associated with empathy, did not differ strongly enough to lead to significant differences between the commander position and the intermediary position. However, in Study 2 using electroencephalography, we observed that responses in EEG potentials that discriminate Shock from NoShock observation were higher for commanders than for intermediaries, but only when commanders were giving orders to a robot. Giving orders to an entity which does not have its own individual responsibility is likely to prevent a diffusion of responsibility phenomenon (Bandura, 2006). This effect was more reliable over the P3 than over the eLPP and the lLPP, which is consistent with former studies (Pech & Caspar, under review; Galang et al., 2021). To investigate further which areas mediate this difference we performed source reconstruction on our EEG data, which revealed involvement of insula and ACC as we had initially hypothesized. An obvious explanation for the difference between the MRI and EEG results in the insula and ACC, is that corrections for multiple comparisons in MRI required a much stricter statistical criterion (p<0.001) compared to the EEG analysis. Indeed our MRI study had a statistical power of 80% only for detecting effects of at least d=0.9. None of the significant effects we observed in the EEG had such large effect sizes. Indeed when looking at uncorrected fMRI results for the CommanderOfRobotAgent(S-NS)-IntermediaryWithHumanAgent(S-NS) we observed ACC activation and when looking at uncorrected results for CommanderOfRobotAgent(S-NS)-CommanderOfHumanAgent(S-NS) we observed ACC and insula activation. Another possible explanation on the difference between the MRI results in Study 1, which did not show statistical difference between our experimental conditions, and the EEG results in Study 2, which showed a higher amplitude of the P3 when participants commanded a robot could be that action-outcome have a shorter delays in EEG than in MRI set-ups. Indeed, in MRI, the outcome followed the agent’s keypress by 3-9s, while in EEG by 200-800ms, because of the long intervals that are needed in fMRI task design. Former studies indicated that action-outcome delays impact agency, with longer action-outcome intervals impacting the sense of agency (Humphreys & Buehner, 2009). It could be the case that empathy is also impacted by long action-outcome delays.

The overlap between the MRI results in Study 1 and the source reconstruction result in EEG in Study 2 showed convergences but also differences. We indeed observed involvement of the AAC and the insula in our EEG data, which is consistent with the MRI results on pain observation in Study 1. However, there were also some areas which were not overlapping. A first critical difference was the auditory tone present after each keypress in the EEG study but not in the MRI study. A second one is that in the EEG study, the shock appeared 200, 500 or 800 ms after the keypress, thus including a lower separation between the motor response and the pain response compared to the MRI study where we use intervals of minimum 3s between the keypress and the shock. A third difference is that in the EEG study, we also asked our participants to estimate the delay between the keypress and the tone, a cognitive task that was not present in the MRI study. Future studies where EEG is used within MRI on the same participants could reveal a more precise overlapping in this context.

When comparing activiations from Study 1 to the activations obtained from another MRI study with the same experimental set-up but participant having the position of the agent (Caspar, Ioumpa et al., 2020), we observed that neural activations in areas including IPL and fusiform gyrus were reduced when individuals are commanding another agent compared to when they are agents themselves. In other words, being free to decide which orders to ask another person to execute leads to a more reduced activation in social cognition related brain regions than being free to both deciding and acting.

When we compared activation patterns between agents coerced and commanders giving orders freely - thus having a classical hierarchical chain between one giving orders and one obeying orders -, the agent had higher brain activation than the commander in empathy related areas as SII and IPL, suggesting that acting has a higher influence of the neural empathic response than having decisional power. The neurological signature results are also supporting this notion. These findings suggest that, while having a low social power reduces the neural empathic response regarding, not being the author of the action leading to the outcome, impacts even more the neural empathic responses.

We also observed that behavioral results slightly differ between study 1 and study 2. In Study 1, the number of shocks delivered to the victim did not statistically change across the three experimental conditions, while in Study 2, agents delivered less shocks in the CommanderOfHumanAgent condition and in the CommanderOfRobotAgent condition, compared to the IntermediaryWithAHumanAgent condition. We actually observed that participants tended to disobey slightly more the orders of the experimenter in the MRI study compared to the EEG study. A possible explanation is that, even though participants were performing the task with the experimenter close to them in the MRI scanner, social distance with the experimenter could have been perceived as higher due to the MRI scanner and headphones. Also, we observed that in Study 1, participants reported a higher feeling of responsibility in both the CommanderOfHumanAgent condition and the CommanderOfRobotAgent condition compared to the IntermediaryWithAHumanAgent condition, while in Study 2, participants reported more responsibility in the CommanderOfRobotAgent condition compared to the two other conditions. This later finding perhaps explain why we also observed a higher neural response to the pain of the other in the CommanderOfRobotAgent condition, as previous studies showed a position relationship between the feeling of responsibility and empathy for pain (Lepron et al., 2015; Cui et al., 2012).

In hierarchical situations, one person decides and orders, and another person executes. Thus, deciding and acting are two different cognitive functions that are split across two different individual’s brains. The data acquired in the present study combined with the data collected in a former study (Caspar, Ioumpa et al., 2020) suggest that being the commander or the intermediary involved reduced brain activations in empathy-related brain regions for the pain inflicted for the victim compared to being free agents that can decide and act themselves. Our results also suggest that neither coerced agents or commanders experience agency over their actions and its consequences. These results show how powerful hierarchical situations can facilitate the commission of actions that harm others, as agency and empathy are split across multiple individuals.

## Author contributions

E.A.C. developed the study concept. E.A.C., K.I., C.K., and V.G. contributed to the study design. Testing and data collection were performed by E.A.C., K.I. and I.A. for Study 1 and E.A.C. for Study 2. E.A.C., K.I., I.A. and L.D.A. performed the data analysis and interpretation under the supervision of C.K. and V.G. E.A.C. and K.I. drafted the manuscript, and C.K., and V.G. provided critical revisions. All authors approved the final version of the manuscript for submission.

## Acknowledgements

The research was funded by the European Union’s Horizon 2020 research and innovation programme under the Marie Skłodowska-Curie grant agreement Agent No 743685 to E.A.C., and from the Netherlands Organization for Scientific Research (VICI: 453–15–009 to C.K. and VIDI 452–14–015 to V.G.).

## REFERENCES

Altemeyer, B. (1981). Right Wing Authoritarianism.Winnipeg: University of Manitoba Press. https://doi.org/10.1038/nrn.2017.14

Angst, J., Adolfsson, R., Benazzi, F., Gamma, A., Hantouche, E., Meyer, T. D., Skeppar, P., Vieta, E., & Scott, J. (2005). The HCL-32: Towards a self-assessment tool for hypomanic symptoms in outpatients. Journal of Affective Disorders, 88(2), 217–233. https://doi.org/10.1016/j.jad.2005.05.011

Ashburner, J., Barnes, G., Chen, C.-C., Daunizeau, J., Flandin, G., Friston, K., Kiebel, S., Kilner, J., Litvak, V., & Moran, R. (2014). SPM12 manual. Wellcome Trust Centre for Neuroimaging, London, UK, 2464.

Balconi, M. (2010). The Sense of Agency in Psychology and Neuropsychology. In B. Michela (Ed.), Neuropsychology of the Sense of Agency: From Consciousness to Action (pp. 3–22). Springer Milan. https://doi.org/10.1007/978-88-470-1587-6_1

Bandura, A. (2006). Toward a Psychology of Human Agency. Perspectives on Psychological Science, 1(2), 164–180. https://doi.org/10.1111/j.1745-6916.2006.00011.x

Bandura, A., Freeman, W. H., & Lightsey, R. (1999). Self-Efficacy: The Exercise of Control. Journal of Cognitive Psychotherapy, 13(2), 158–166. https://doi.org/10.1891/0889-8391.13.2.158

Barlas, Z., & Obhi, S. (2013). Freedom, choice, and the sense of agency. Frontiers in Human Neuroscience, 7, 514. https://doi.org/10.3389/fnhum.2013.00514

Benjamini, Y., & Hochberg, Y. (1995). Controlling the false discovery rate: A practical and powerful approach to multiple testing. Journal of the Royal Statistical Society B, 57(1), 289–300. https://doi.org/10.2307/2346101

Bernhardt, B. C., & Singer, T. (2012). The Neural Basis of Empathy. Annual Review of Neuroscience, 35(1), 1–23. https://doi.org/10.1146/annurev-neuro-062111-150536

Buehner, M. J., & Humphreys, G. R. (2009). Causal Binding of Actions to Their Effects. Psychological Science, 20(10), 1221–1228. https://doi.org/10.1111/j.1467-9280.2009.02435.x

Caspar, E. A., Christensen, J. F., Cleeremans, A., & Haggard, P. (2016). Coercion Changes the Sense of Agency in the Human Brain. Current Biology, 26(5), 585–592. https://doi.org/10.1016/j.cub.2015.12.067

Caspar, E. A., Cleeremans, A., & Haggard, P. (2018). Only giving orders? An experimental study of the sense of agency when giving or receiving commands. PLOS ONE, 13(9), e0204027. https://doi.org/10.1371/journal.pone.0204027

Caspar, E. A., Ioumpa, K., Keysers, C., & Gazzola, V. (2020). Obeying orders reduces vicarious brain activation towards victims’ pain. NeuroImage, 222, 117251. https://doi.org/10.1016/j.neuroimage.2020.117251

Caspar, E. A., Lo Bue, S., Magalhães De Saldanha da Gama, P. A., Haggard, P., & Cleeremans, A. (2020). The effect of military training on the sense of agency and outcome processing. Nature Communications, 11(1), 4366. https://doi.org/10.1038/s41467-020-18152-x

Chen, Y.-C., Chen, C.-C., Decety, J., & Cheng, Y. (2014). Aging is associated with changes in the neural circuits underlying empathy. Neurobiology of Aging, 35(4), 827–836. https://doi.org/10.1016/j.neurobiolaging.2013.10.080

Ciardo, F., Beyer, F., De Tommaso, D., & Wykowska, A. (2020). Attribution of intentional agency towards robots reduces one’s own sense of agency. Cognition, 194, 104109. https://doi.org/10.1016/j.cognition.2019.104109

Coll, M.-P. (2018). Meta-analysis of ERP investigations of pain empathy underlines methodological issues in ERP research. Social Cognitive and Affective Neuroscience, 13(10), 1003–1017. https://doi.org/10.1093/scan/nsy072

Cravo, A. M., Haddad, H., Claessens, P. M. E., & Baldo, M. V. C. (2013). Bias and learning in temporal binding: Intervals between actions and outcomes are compressed by prior bias. Consciousness and Cognition, 22(4), 1174–1180. https://doi.org/10.1016/j.concog.2013.08.001

Cui, F., Abdelgabar, A.-R., Keysers, C., & Gazzola, V. (2015). Responsibility modulates pain-matrix activation elicited by the expressions of others in pain. NeuroImage, 114, 371–378. https://doi.org/10.1016/j.neuroimage.2015.03.034

Dale, A. M., & Sereno, M. I. (1993). Improved Localizadon of Cortical Activity by Combining EEG and MEG with MRI Cortical Surface Reconstruction: A Linear Approach. Journal of Cognitive Neuroscience, 5(2), 162–176. https://doi.org/10.1162/jocn.1993.5.2.162

Davis, M. H., & Association, A. P. (1980). A multidimensional approach to individual differences in empathy. JSAS Catalog of Selected Documents in Psychology.

Decety, J. (2011). Dissecting the Neural Mechanisms Mediating Empathy. Emotion Review, 3(1), 92–108. https://doi.org/10.1177/1754073910374662

Dienes, Z. (2011). Bayesian Versus Orthodox Statistics: Which Side Are You On? Perspectives on Psychological Science, 6(3), 274–290. https://doi.org/10.1177/1745691611406920

Dunwoody, P. T., & Funke, F. (2016). The Aggression-Submission-Conventionalism Scale: Testing a New Three Factor Measure of Authoritarianism. Journal of Social and Political Psychology, 4(2), 571–600. https://doi.org/10.5964/jspp.v4i2.168

Eickhoff, S. B., Stephan, K. E., Mohlberg, H., Grefkes, C., Fink, G. R., Amunts, K., & Zilles, K. (2005). A new SPM toolbox for combining probabilistic cytoarchitectonic maps and functional imaging data. Neuroimage, 25(4), 1325–1335.

Galang, C. M., Jenkins, M., Fahim, G., & Obhi, S. S. (2021). Exploring the relationship between social power and the ERP components of empathy for pain. Social Neuroscience, 16(2), 174–188. https://doi.org/10.1080/17470919.2021.1886165

Gallagher, H. L., Jack, A. I., Roepstorff, A., & Frith, C. D. (2002). Imaging the Intentional Stance in a Competitive Game. NeuroImage, 16(3, Part A), 814–821. https://doi.org/10.1006/nimg.2002.1117

Gallagher, S. (2000). Philosophical conceptions of the self: Implications for cognitive science. Trends in Cognitive Sciences, 4(1), 14–21. https://doi.org/10.1016/S1364-6613(99)01417-5

Gallo, S., Paracampo, R., Müller-Pinzler, L., Severo, M. C., Blömer, L., Fernandes-Henriques, C., Henschel, A., Lammes, B. K., Maskaljunas, T., Suttrup, J., Avenanti, A., Keysers, C., & Gazzola, V. (2018). The causal role of the somatosensory cortex in prosocial behaviour. ELife, 7. https://doi.org/10.7554/eLife.32740

Geday, J., Gjedde, A., Boldsen, A.-S., & Kupers, R. (2003). Emotional valence modulates activity in the posterior fusiform gyrus and inferior medial prefrontal cortex in social perception. NeuroImage, 18(3), 675–684. https://doi.org/10.1016/S1053-8119(02)00038-1

Graham, J., Nosek, B. A., Haidt, J., Iyer, R., Koleva, S., & Ditto, P. H. (2011). Mapping the moral domain. Journal of Personality and Social Psychology, 101(2), 366–385. https://doi.org/10.1037/a0021847

Haggard, P. (2017). Sense of agency in the human brain. Nature Reviews Neuroscience, 18(4), 196–207. https://doi.org/10.1038/nrn.2017.14

Humphreys, G. R., & Buehner, M. J. (2009). Magnitude estimation reveals temporal binding at super-second intervals. Journal of Experimental Psychology: Human Perception and Performance, 35(5), 1542. https://doi.org/10.18637/jss.v088.i02

Luck, S. J., & Kappenman, E. S. (Eds.). (2011). The Oxford handbook of event-related potential components. Oxford university press.

Janowski, V., Camerer, C., & Rangel, A. (2013). Empathic choice involves vmPFC value signals that are modulated by social processing implemented in IPL. Social Cognitive and Affective Neuroscience, 8(2), 201–208. https://doi.org/10.1093/scan/nsr086

Jauniaux, J., Khatibi, A., Rainville, P., & Jackson, P. L. (2019). A meta-analysis of neuroimaging studies on pain empathy: Investigating the role of visual information and observers’ perspective. Social Cognitive and Affective Neuroscience, 14(8), 789–813. https://doi.org/10.1093/scan/nsz055

Keysers, C., & Gazzola, V. (2014). Dissociating the ability and propensity for empathy. Trends in Cognitive Sciences, 18(4), 163–166. https://doi.org/10.1016/j.tics.2013.12.011

Keysers, C., Gazzola, V., & Wagenmakers, E.-J. (2020). Using Bayes factor hypothesis testing in neuroscience to establish evidence of absence. Nature Neuroscience, 23(7), 788–799. https://doi.org/10.1038/s41593-020-0660-4

Kilham, W., & Mann, L. (1974). Level of destructive obedience as a function of transmitter and executant roles in the Milgram obedience paradigm. Journal of Personality and Social Psychology, 29(5), 696–702. https://doi.org/10.1037/h0036636

Lamm, C., Decety, J., & Singer, T. (2011). Meta-analytic evidence for common and distinct neural networks associated with directly experienced pain and empathy for pain. NeuroImage, 54(3), 2492–2502. https://doi.org/10.1016/j.neuroimage.2010.10.014

Lamm, C., Rütgen, M., & Wagner, I. C. (2019). Imaging empathy and prosocial emotions. Neuroscience Letters, 693, 49–53. https://doi.org/10.1016/j.neulet.2017.06.054

Lepron, E., Causse, M., & Farrer, C. (2015). Responsibility and the sense of agency enhance empathy for pain. Proceedings of the Royal Society B: Biological Sciences, 282(1799), 20142288. https://doi.org/10.1098/rspb.2014.2288

Levenson, M. R., Kiehl, K. A., & Fitzpatrick, C. M. (1995). Assessing psychopathic attributes in a noninstitutionalized population. Journal of Personality and Social Psychology, 68(1), 151–158. https://doi.org/10.1037/0022-3514.68.1.151

Li, P., Han, C., Lei, Y., Holroyd, C. B., & Li, H. (2011). Responsibility modulates neural mechanisms of outcome processing: An ERP study. Psychophysiology, 48(8), 1129–1133. https://doi.org/10.1111/j.1469-8986.2011.01182.x

Lockwood, P. L., Sebastian, C. L., McCrory, E. J., Hyde, Z. H., Gu, X., De Brito, S. A., & Viding, E. (2013). Association of callous traits with reduced neural response to others’ pain in children with conduct problems. Current Biology, 23(10), 901–905. https://doi.org/10.1016/j.cub.2013.04.018

Love, J., Selker, R., Marsman, M., Jamil, T., Dropmann, D., Verhagen, A. J., Ly, A., Gronau, Q. F., Šmíra, M., Epskamp, S., Matzke, D., Wild, A., Knight, P., Rouder, J. N., Morey, R. D., Wagenmakers, E. J. E.-J., Verhagen, J., Ly, A., Gronau, Q. F., … Wagenmakers, E. J. E.-J. (2019). JASP: Graphical statistical software for common statistical designs. Journal of Statistical Software, 88(2), 1–17. https://doi.org/10.18637/jss.v088.i02

Luck, S. J. (2012). Electrophysiological correlates of the focusing of attention within complex visual scenes: N2pc and related ERP components. In The Oxford handbook of event-related potential components (pp. 329–360). Oxford University Press.

Meck, W. H. (2006). Neuroanatomical localization of an internal clock: A functional link between mesolimbic, nigrostriatal, and mesocortical dopaminergic systems. Brain Research, 1109(1), 93–107. https://doi.org/10.1016/j.brainres.2006.06.031

Milgram, S. (1974). The Dilemma of Obedience. The Phi Delta Kappan, 55(9), 603–606.

Obhi, S. S., Swiderski, K. M., & Brubacher, S. P. (2012). Induced power changes the sense of agency. Consciousness and Cognition, 21(3), 1547–1550. https://doi.org/10.1016/j.concog.2012.06.008

Oostenveld, R., Fries, P., Maris, E., & Schoffelen, J.-M. (2011). FieldTrip: Open source software for advanced analysis of MEG, EEG, and invasive electrophysiological data. Computational Intelligence and Neuroscience, 2011, 156869. https://doi.org/10.1155/2011/156869

Poldrack, R. A. (2006). Can cognitive processes be inferred from neuroimaging data? Trends in Cognitive Sciences, 10(2), 59–63. https://doi.org/10.1016/j.tics.2005.12.004

Rouder, J. N., Morey, R. D., Speckman, P. L., & Province, J. M. (2012). Default Bayes factors for ANOVA designs. Journal of Mathematical Psychology, 56(5), 356–374. https://doi.org/10.1016/j.jmp.2012.08.001

Schulte-Rüther, M., Markowitsch, H. J., Fink, G. R., & Piefke, M. (2007). Mirror Neuron and Theory of Mind Mechanisms Involved in Face-to-Face Interactions: A Functional Magnetic Resonance Imaging Approach to Empathy. Journal of Cognitive Neuroscience, 19(8), 1354–1372. https://doi.org/10.1162/jocn.2007.19.8.1354

Swaan, A. de. (2015). The Killing Compartments: The Mentality of Mass Murder. In The Killing Compartments. Yale University Press. https://doi.org/10.12987/9780300210675

Timmers, I., Park, A. L., Fischer, M. D., Kronman, C. A., Heathcote, L. C., Hernandez, J. M., & Simons, L. E. (2018). Is Empathy for Pain Unique in Its Neural Correlates? A Meta-Analysis of Neuroimaging Studies of Empathy. Frontiers in Behavioral Neuroscience, 12. https://doi.org/10.3389/fnbeh.2018.00289

Wagenmakers, E.-J., Verhagen, A. J., & Ly, A. (2016). How to Quantify the Evidence for the Absence of a Correlation. Behavior Research Methods, 48, 413–426.

Zaki, J. (2014). Empathy: A motivated account. Psychological Bulletin, 140(6), 1608–1647. https://doi.org/10.1037/a0037679

Zaki, J., Bolger, N., & Ochsner, K. (2009). Unpacking the Informational Bases of Empathic Accuracy. Emotion, 9(4), 478–487. https://doi.org/10.1037/a0016551

Zhou, F., Li, J., Zhao, W., Xu, L., Zheng, X., Fu, M., Yao, S., Kendrick, K. M., Wager, T. D., & Becker, B. (2020). Empathic pain evoked by sensory and emotional-communicative cues share common and process-specific neural representations. ELife, 9, e56929. https://doi.org/10.7554/eLife.56929

