## SupplementaryInformation for "Commanding or being a simple intermediary: how does it affect moral behavior and related brain mechanisms?"

### *SUPPLEMENTARY INFORMATION S1*

**Supplementary Table 1. BOLD activity of Shocks - Noshock contrast for all experimental conditions together.** Only clusters surviving a 5% FWE correction at the cluster size are reported (t=3.5, p < .001, cluster size 160). Brain regions are identified using the Anatomy Toolbox (Eickhoff et al., 2005).

| Cluster size | Voxels in cyto | % Cluster | Hem | Cyto or Anatomical description | % Area | Peak t-value | MNI coordinates | | |
|---|---|---|---|---|---|---|---|---|---|
| | | | | | | | x | y | z |
| | | | | **Shocks - Noshock all experimental conditions** (5% FWE correction . t=3.5 . p<.001 . k=160) | | | | | |
| 3622 | | | | Area hOc4la | | | | | |
| | 522.9 | 14.4 | L | Middle Temporal Gyrus | 61.2 | 10.49 | -54 | -70 | 2 |
| | | | | Area hOc4lp | | | | | |
| | 460.6 | 12.7 | L | Middle Occipital Gyrus | 53.8 | 5.98 | -28 | -92 | 12 |
| | | | | Area hOc4v [V4(v)] | | | | | |
| | 262.5 | 7.2 | L | Inferior Occipital Gyrus | 36.1 | 7.07 | -30 | -82 | -6 |
| | | | | Area hOc4v [V4(v)] | | | | | |
| | | | L | Lingual Gyrus | | 5.71 | -32 | -86 | -14 |
| | | | | Lobule VIIa crusI | | | | | |
| | 231.1 | 6.4 | L | Cerebelum | 7.6 | 5.11 | -26 | -66 | -32 |
| | | | | Area FG2 | | | | | |
| | 192.8 | 5.3 | L | Fusiform Gyrus | 37.8 | 8.04 | -42 | -68 | -14 |
| | | | | Area FG3 | | | | | |
| | 183.4 | 5.1 | L | Inferior Temporal Gyrus | 22.2 | 6.30 | -40 | -42 | -18 |
| | | | | Area FG3 | | | | | |
| | | | L | Fusiform Gyrus | | 5.88 | -40 | -54 | -20 |
| | 143.4 | 4 | L | Area hOc3v [V3v] | 15.5 | | | | |
| | 131.3 | 3.6 | L | Lobule VI | 7 | | | | |
| | | | | Area hOc5 [V5/MT] | | | | | |
| | 81.5 | 2.3 | L | Inferior Occipital Gyrus | 101.4 | 8.73 | -40 | -72 | -4 |
| | 70.1 | 1.9 | L | Area hOc3d [V3d] | 7.1 | | | | |
| | 67.5 | 1.9 | L | Area hOc1 [V1] | 3.3 | | | | |
| | 20.6 | 0.6 | L | Area FG1 | 8.1 | | | | |
| | 18.6 | 0.5 | L | Area hOc2 [V2] | 2 | | | | |
| | 11.4 | 0.3 | L | Area hOc4d [V3A] | 2 | | | | |
| | 4.9 | 0.1 | L | Area FG4 | 0.8 | | | | |
| | 3 | 0.1 | L | Area PGp (IPL) | 0.4 | | | | |
| 3352 | | | | Area 45 | | | | | |
| | 448.4 | 448.4 | R | R IFG (p. Opercularis) | 43.4 | 7.29 | 56 | 18 | 32 |
| | | | | Area 45 | | 6.81 | | | |
| | | | R | IFG (p. Triangularis) | | | 52 | 20 | 24 |
| | | | | Area 44 | | | | | |
| | 256.5 | 256.5 | R | R IFG (p. Opercularis) | 42.7 | 7.88 | 58 | 16 | 6 |
| | 13 | 13 | R | Area Fo2 | 1.2 | | | | |
| | 8 | 8 | R | Area Id1 | 4.9 | | | | |
| | 1.4 | 1.4 | R | Thal: Parietal | 0.4 | | | | |
| | 0.8 | 0.8 | R | Thal: Temporal | 0.1 | | | | |
| | | | R | Insula Lobe | | 9.33 | 38 | 20 | -2 |
| | | | R | Insula Lobe | | 6.30 | 28 | 20 | -16 |
| | | | R | Middle Frontal Gyrus | | 7.25 | 46 | 48 | 4 |
| | | | R | Precentral Gyrus | | 6 | 46 | 6 | 36 |
| 2850 | 401.6 | 14.1 | R | Area hOc4la | 45.3 | | | | |
| | | | | Area hOc4lp | | | | | |
| | 252 | 8.8 | R | Middle Temporal Gyrus | 45 | 8.16 | 44 | -72 | 2 |
| | | | | Area hOc4lp | | | | | |
| | | | R | Middle Occipital Gyrus | | 6.69 | 46 | -78 | 6 |
| | 202.1 | 7.1 | R | Lobule VI (Hem) | 11.2 | | | | |
| | 181.8 | 6.4 | R | Lobule VIIa crusI (Hem) | 5.6 | | | | |
| | | | | Area FG3 | | | | | |
| | 112 | 3.9 | R | Inferior Temporal Gyrus | 17.1 | 6.66 | 42 | -56 | -14 |
| | 81 | 2.8 | R | Area hOc4v [V4(v)] | 13 | | | | |
| | 79.5 | 2.8 | R | Area FG2 | 24.4 | | | | |
| | | | | Area hOc5 [V5/MT] | | | | | |
| | 56.8 | 2 | R | Middle Temporal Gyrus | 97.4 | 8.91 | 46 | -66 | 4 |
| | 44.9 | 1.6 | R | Area hOc3v [V3v] | 5.3 | | | | |

| | | | | | | | | | |
|---|---|---|---|---|---|---|---|---|---|
| | 31.9 | 1.1 | R | Area FG1 | 12.8 | | | | |
| | | | R | Area hOc3d [V3d] Middle | | | | | |
| | 21.3 | 0.7 | | Occipital Gyrus | 3.9 | 5.94 | 30 | -94 | 6 |
| | 11.9 | 0.4 | R | Area hOc4d [V3A] | 2.8 | | | | |
| | 10.5 | 0.4 | R | Lobule VI (Verm) | 4.5 | | | | |
| | 5.9 | 0.2 | R | Area hOc1 [V1] | 0.3 | | | | |
| | 5.9 | 0.2 | R | Area FG4 | 1.2 | | | | |
| | 5.6 | 0.2 | R | Area PGp (IPL) | 0.6 | | | | |
| | 3.5 | 0.1 | R | Area PGa (IPL) | 0.5 | | | | |
| | | | R | Inferior Temporal Gyrus | | 9.15 | 44 | -68 | -4 |
| 1854 | 371 | 20 | L | Thal: Prefrontal | 58.8 | 6.48 | 4 | -6 | 2 |
| | 303.5 | 16.4 | R | Thal: Prefrontal | 54.2 | 5.74 | 14 | -2 | 6 |
| | | | R | Thal: Prefrontal | | | | | |
| | | | | Thalamus | | 5.54 | 10 | -8 | 6 |
| | 142.5 | 7.7 | R | Thal: Temporal | 26.1 | 5.92 | -12 | -8 | 6 |
| | | | R | Thal: Temporal | | | | | |
| | | | | Thalamus | | 5.50 | 8 | -14 | 10 |
| | 93.4 | 5 | L | Thal: Temporal | 17.6 | 5.54 | -2 | -8 | 2 |
| | 15.3 | 0.8 | L | Thal: Premotor | 12.9 | | | | |
| | 1.4 | 0.1 | R | Thal: Parietal | 0.4 | | | | |
| | 0.3 | 0 | L | Thal: Parietal | 0.1 | | | | |
| | | | L | Pallidum | | 5.97 | -12 | 2 | -2 |
| 1227 | | | R | Area hIP3 (IPS) | | | | | |
| | 252 | 20.5 | | Inferior Parietal Lobule | 55.2 | 5.36 | 34 | -54 | 48 |
| | | | R | Area PF (IPL) | | 5.30 | | | |
| | 154 | 12.6 | | SupraMarginal Gyrus | 22.8 | | 64 | -32 | 24 |
| | | | R | Area PFm (IPL) | | | | | |
| | 153.1 | 12.5 | | SupraMarginal Gyrus | 21.7 | 6.12 | 54 | -44 | 42 |
| | | | R | Area PFcm (IPL) Superior | | | | | |
| | 107.6 | 8.8 | | Temporal Gyrus | 33 | 5.30 | 60 | -36 | 20 |
| | | | R | Area 7PC (SPL) | | 5.95 | | | |
| | 89.1 | 7.3 | | Superior Parietal Lobule | 19.6 | | 34 | -54 | 60 |
| | | | R | Area 7PC (SPL) | | | | | |
| | | | | Inferior Parietal Lobule | | 4.89 | 32 | -46 | 50 |
| | 71 | 5.8 | R | Area PFt (IPL) | 17 | | | | |
| | | | R | Area PFop (IPL) | | | | | |
| | 57.6 | 4.7 | | SupraMarginal Gyrus | 25.2 | 4.55 | 58 | -20 | 26 |
| | 48.1 | 3.9 | R | Area 2 | 7.4 | | | | |
| | 41.9 | 3.4 | R | Area hIP2 (IPS) | 19.9 | | | | |
| | 26.1 | 2.1 | R | Area hIP1 (IPS) | 9 | | | | |
| | 22.6 | 1.8 | R | Area 7A (SPL) | 2.9 | | | | |
| | 4.5 | 0.4 | R | Area PGa (IPL) | 0.6 | | | | |
| | 0.9 | 0.1 | R | Area 1 | 0.1 | | | | |
| | 0.6 | 0.1 | R | Area OP1 [SII] | 0.2 | | | | |
| | 0.6 | 0.1 | R | Area 3b | 0.1 | | | | |
| | 0.3 | 0 | R | Area 3a | 0.1 | | | | |
| 947 | | | L | Superior Medial Gyrus | | 6.98 | 2 | 32 | 50 |
| | | | R | ACC | | 6.70 | 8 | 34 | 20 |
| | | | R | Posterior-Medial Frontal | | 6.44 | 4 | 22 | 58 |
| | | | R | Superior Medial Gyrus | | 6.44 | 4 | 30 | 42 |
| | | | L | ACC | | 4.95 | 0 | 30 | 30 |
| 933 | | | L | PCC | | 6.15 | -8 | -42 | 24 |
| 705 | | | L | Insula | | 8.98 | -30 | 18 | -10 |
| | | | L | IFG (p. Triangularis) | | 5.89 | -42 | 20 | 8 |
| 278 | | | L | Area hIP3 (IPS) | | | | | |
| | 118 | 42.4 | | Inferior Parietal Lobule | 25.8 | 5.72 | -34 | -46 | 48 |
| | 26.3 | 9.4 | L | Area hIP1 (IPS) | 7.2 | | | | |
| | 18.5 | 6.7 | L | Area 7PC (SPL) | 10.8 | | | | |
| | | | L | Area 7A (SPL) | | | | | |
| | 1.8 | 0.6 | | Superior Parietal Lobule | 0.1 | 3.81 | -30 | -56 | 56 |
| | 1.1 | 0.4 | L | Area 5L (SPL) | 0.2 | | | | |
| | 0.5 | 0.2 | L | Area 2 | 0.1 | | | | |
| 162 | | | L | Precuneus | | 4.65 | -10 | -60 | 32 |
| 160 | | | R | Middle Temporal Gyrus | | 5.06 | 56 | -30 | -8 |

### *SUPPLEMENTARY INFORMATION S2*

**Supplementary Table 2. BOLD activity of comparisons between the agent and commander study conditions.** Only clusters surviving a 5% FWE correction at the cluster size are reported (t=3.5, p < .001, cluster size 160). Brain regions are identified using the Anatomy Toolbox (Eickhoff et al. 2005).

| Cluster size | Voxels in cyto | % Cluster | Hem | Cyto or Anatomical description | % Area | Peak t-value | MNI coordinates | | |
|---|---|---|---|---|---|---|---|---|---|
| | | | | | | | x | y | z |
| **Two sample t-tests between Agent Free Shocks - Noshock and Intermediary with Human Agent Shocks - Noshock** (5% FWE correction . t=3.5 . p<.001 . k=236) | | | | | | | | | |
| 373 | 97 | 26 | L | Area PFt (IPL) Postcentral Gyrus | 16.6 | 4.43 | -52 | -22 | 28 |
| | | | L | Area PFt (IPL) SupraMarginal Gyrus | | 3.59 | -62 | -26 | 32 |
| | 95.6 | 25.6 | L | Area PFop (IPL) Postcentral Gyrus | 43.1 | 4.48 | -58 | -20 | 26 |
| | 66.3 | 17.8 | L | Area PF (IPL) SupraMarginal Gyrus | 12.7 | 4.74 | -64 | -32 | 28 |
| | 27 | 7.2 | L | Area PFcm (IPL) | 8.3 | | | | |
| | 15.4 | 4.1 | L | Area 3b Postcentral Gyrus | 2.7 | 4.37 | -48 | -22 | 32 |
| | 12 | 3.2 | L | Area 2 | 2.3 | | | | |
| | 4.9 | 1.3 | L | Area OP1 [SII] | 1.3 | | | | |
| | 4.1 | 1.1 | L | Area 3a | 1.4 | | | | |
| | 1.8 | 0.5 | L | Area OP4 [PV] | 0.5 | | | | |
| | 1.1 | 0.3 | L | Area TE 3 | 0.1 | | | | |
| | 0.5 | 0.1 | L | Area 1 | 0.1 | | | | |
| | | | L | Superior Temporal Gyrus | | 3.51 | -62 | -40 | 14 |
| 237 | 50.3 | 21.2 | R | Area PFcm (IPL) Superior Temporal Gyrus | 15.4 | 3.63 | 60 | -32 | 18 |
| | 39 | 16.5 | R | Area PF (IPL) Superior Temporal Gyrus | 5.8 | 4.48 | 60 | -40 | 18 |
| | 37.5 | 15.8 | R | Area PFm (IPL) | 5.3 | | | | |
| | 7.9 | 3.3 | R | Area PGa (IPL) | 1.1 | | | | |
| | | | R | Middle Temporal Gyrus | | 3.66 | 60 | -46 | 10 |
| 236 | 7.5 | 3.2 | R | Area 44 | 1.2 | | | | |
| | 0.4 | 0.2 | R | Area 45 | 0 | | | | |
| | | | R | IFG (p. Opercularis) | | 4.81 | 46 | 8 | 28 |
| | | | R | Precentral Gyrus | | 4.03 | 48 | 6 | 42 |
| **Two sample t-tests between Agent Free Shocks - Noshock and Commander of Human Agent Shocks - Noshock** (5% FWE correction . t=3.5 . p<.001 . k=163) | | | | | | | | | |
| 258 | 33.3 | 12.9 | R | Area FG4 Fusiform Gyrus | 6.8 | 4.48 | 30 | -52 | -10 |
| | 5 | 1.9 | R | Lobule V (Hem) | 0.6 | | | | |
| | 3.4 | 1.3 | R | Subiculum | 0.9 | | | | |
| | 2.6 | 1 | R | Area FG1 | 1.1 | | | | |
| | 0.9 | 0.3 | R | Area hOc3v [V3v] | 0.1 | | | | |
| | 0.4 | 0.1 | R | Area hOc1 [V1] | 0 | | | | |
| | 0.3 | 0.1 | R | Area hOc2 [V2] | 0 | | | | |
| | | | R | Fusiform Gyrus | | 4.86 | 26 | -42 | -12 |
| | | | R | ParaHippocampal Gyrus | | 3.90 | 18 | -42 | -6 |
| | | | R | Cerebelum (IV-V) | | 3.80 | 22 | -36 | -20 |
| 232 | 75.9 | 32.7 | L | Area FG4 Fusiform Gyrus | 12.8 | 5.59 | -26 | -52 | -10 |
| | 33 | 14.2 | L | Area hOc1 [V1] | 1.6 | | -14 | -58 | -4 |

| | | | | | | | | | |
|---|---|---|---|---|---|---|---|---|---|
| | | | | Lingual Gyrus | | 4.04 | | | |
| | 10.3 | 4.4 | L | Area hOc2 [V2] | 1.1 | | | | |
| | 5 | 2.2 | L | Area hOc3v [V3v] | 0.5 | | | | |
| | 1.1 | 0.5 | L | Area hOc4v [V4(v)] | 0.2 | | | | |
| 163 | 23 | 14.1 | R | Area PF (IPL) | 3.4 | | | | |
| | | | R | Area PFm (IPL) | | | | | |
| | 22 | 13.5 | | Superior Temporal Gyrus | 3.1 | 4.20 | 62 | -42 | 20 |
| | 7 | 4.3 | R | Area PGa (IPL) | 0.9 | | | | |
| | 6.6 | 4.1 | R | Area PFcm (IPL) | 2 | | | | |
| | | | R | Superior Temporal Gyrus | | 4.02 | 60 | -46 | 14 |
| | | | R | Middle Temporal Gyrus | | 3.98 | 60 | -44 | 10 |
| **Two sample t-tests between Agent Coerced Shocks - Noshock and Commander of Human Agent Shocks - Noshock** (5% FWE correction . t=3.5 . p<.001 . k=315) | | | | | | | | | |
| 315 | | | R | Area 4a | | | | | |
| | 37 | 11.7 | | Precentral Gyrus | 3.4 | 4.12 | 22 | -30 | 66 |
| | | | L | Area 4a | | | | | |
| | 7.2 | 2.3 | | Paracentral Lobule | 0.8 | 3.77 | -4 | -24 | 70 |
| | 4.7 | 1.5 | R | Area 4p | 1.5 | | | | |
| | | | R | Posterior-Medial Frontal | | 5.22 | 12 | -20 | 66 |
| | | | R | Postcentral Gyrus | | 4.69 | 12 | -30 | 62 |
| | | | R | Superior Frontal Gyrus | | 4.08 | 18 | -10 | 66 |
